# Discovery and therapeutic exploitation of Master Regulatory miRNAs in Glioblastoma

**DOI:** 10.1101/2025.04.01.646663

**Authors:** Shekhar Saha, Ying Zhang, Myron K Gilbert, Collin Dube, Farina Hanif, Elizabeth Qian Xu Mulcahy, Sylwia Bednarek, Pawel Marcinkiewicz, Xiantao Wang, Gijung Kwak, Kadie Hudson, Yunan Sun, Manikarna Dinda, Tapas Saha, Fadila Guessous, Nichola Cruickshanks, Rossymar Rivera Colon, Lily Dell’Olio, Rajitha Anbu, Benjamin Kefas, Pankaj Kumar, Alexander L Klibanov, David Schiff, Jung Soo Suk, Justin Hanes, Jamie Mata, Markus Hafner, Roger Abounader

## Abstract

Glioblastoma is a fatal primary malignant brain tumor, with an average survival of only 15 months despite surgical resection, chemotherapy, and radiation therapy. Due to the concurrent deregulation of numerous genes in glioblastoma, molecular monotherapies have not improved clinical outcomes. Evidence suggests that effectively targeting multiple deregulated molecules is essential for better therapies; however, this is limited by the lack of suitable drugs and the increased toxicity of combination therapies. To address this, we hypothesized that miRNAs, small gene-regulatory RNAs that suppress multiple target genes via sequence complementarity, could be developed to inhibit multiple deregulated genes simultaneously, leading to more effective treatments. We identified master regulatory miRNAs—those that target several deregulated genes in glioblastoma—using PAR-CLIP screenings in glioblastoma cells and analyzed TCGA tumor data to find which targets were deregulated. An algorithm ranked these targets based on their significance in glioblastoma malignancy. We selected two tumor suppressor master regulatory miRNAs, miR-340 and miR-382, and one oncogenic miRNA, miR-17. Validation showed that these miRNAs target critical glioblastoma pathways and significantly inhibit cell growth, survival, invasion, and tumor growth in vivo. We developed an innovative therapeutic delivery approach using Brain Penetrating Nanoparticles in combination with MRI-guided focused ultrasound and microbubbles, resulting in reduced tumor volume and extended survival in glioblastoma-bearing mice. This strategy offers a promising pathway for translating miRNA-based therapies into clinical trials for glioblastoma and other cancers.

**One Sentence Summary:** We developed and used new computational, experimental, and therapeutic approaches to identify and therapeutically deliver master regulatory miRNAs to inhibit the growth of glioblastoma, the most common and deadly primary brain tumor.

## Introduction

Glioblastoma is an aggressive and fatal primary brain cancer that remains a formidable challenge due to its heterogeneity, therapeutic resistance, and invasive nature(*1–3*). Despite advances in therapies and clinical trials, the standard treatment of maximum surgical resection followed by radiation and temozolomide chemotherapy only extends patient survival to a modest 14.6 months (*4–7*). The Cancer Genome Atlas (TCGA) and other studies comprehensively analyzed deregulated gene expression in several hundred GBM tumors and described the concurrent deregulation of numerous genes in any single tumor (*8, 9*). Because of this multi-gene deregulation, molecular monotherapies have failed to achieve significant improvements in clinical outcomes. Several lines of evidence suggest that simultaneous targeting of several deregulated molecules is required to achieve better therapies (*10–13*). However, the simultaneous targeting of several deregulated oncogenic drivers using conventional drugs is severely limited by the fact that the drugs needed to simultaneously target many deregulated molecules do not currently exist, and because combining several drugs in a clinical setting leads to an exponential increase in toxicity (*14*). The goal of this study was to identify master regulatory microRNAs (miRNAs), defined as miRNAs that target several deregulated genes in glioblastoma, and use them or their inhibitors to simultaneously target multiple deregulated molecules for GBM therapy.

MicroRNAs (miRNAs) are small non-coding RNA molecules that span 19-24 nucleotides. MiRNAs exert their effects by incorporating into argonaute (AGO) proteins (4 AGOs in humans) and guiding them to target mRNA via seed-pairing predominantly to the 3’-untranslated regions (3’-UTR), and less commonly to the coding sequence (CDS), and 5’-untranslated regions (5’-UTR) of target genes. This facilitates the recruitment of effector proteins and the assembly of miRNA-induced silencing complexes (miRISC). The miRISC induces either mRNA degradation or translational inhibition (*15–17*). miRNAs play pivotal roles in regulating various cellular processes, including proliferation, invasion, apoptosis, and differentiation (*18–21*). Dysregulation of miRNA expression is a hallmark of various cancer types, where miRNAs can function as either oncogenes or tumor suppressors by suppressing mRNAs of tumor suppressors or oncogenes, respectively (*22–29*). Importantly, because miRNAs do not require full complementarity to inhibit targeted mRNAs, single miRNAs can target and simultaneously inhibit numerous genes (*30–32*).

We defined “master regulatory miRNAs” as those that target several deregulated genes in glioblastoma. We reasoned that we could identify master regulatory miRNA and then use them for glioblastoma therapy. Using them as therapeutic agents would theoretically be equivalent to using a combination of several drugs that target deregulated glioblastoma driver genes. To find master regulatory miRNAs, we first used PAR-CLIP to identify all targets of all miRNAs in glioblastoma cells (*33*). We then analyzed TCGA tumor data to determine which of these targets are deregulated in human tumors. We developed and implemented a computational algorithm to prioritize miRNA targets based on their relevance to glioblastoma malignancy. We selected the top candidate master regulatory miRNAs, defined by their capacity to regulate numerous target genes and therefore, exhibit strong anti-tumor effects when delivered as therapy.

A major challenge to successful miRNA therapy is delivery. The central nervous system’s protective blood-brain barrier poses a formidable challenge to drug delivery (*34*). Focused Ultrasound (FUS) presents a noninvasive and reversible approach to transiently open the blood-brain barrier (BBB) in animal models (*35*). This temporary BBB opening facilitates drug delivery in brain tumors and other brain diseases (*36, 37*). Recent clinical studies have successfully used Magnetic Resonance Image-guided FUS with microbubbles (MB) to deliver chemotherapeutic drugs to human brains and treat neurodegenerative conditions such as Alzheimer’s disease, Parkinson’s disease, and amyotrophic lateral sclerosis (*38–42*). A second challenge to achieving effective gene therapy distribution in the brain is the extracellular matrix (ECM), a dense and nanoporous network composed of electrostatically charged molecules such as proteoglycans, hyaluronan, and tenascins that restrict the diffusion of gene vectors through steric hindrance and adhesive interactions (*43*). To address this problem, brain-penetrating nanoparticles (BPN) consisting of poly(beta-amino esters) (PBAE) that are coated with a high density of polyethylene glycol (PEG) were developed (*44*).

In this study, we integrated miRNA target identification from PAR-CLIP with gene expression and survival data from TCGA to uncover master regulatory miRNAs in glioblastoma. Through this integrative approach, we identified three key miRNAs that collectively regulate a substantial number of glioblastoma driver genes: miR-340 and miR-382, which act as tumor-suppressive master regulators targeting 87 and 79 genes respectively, and miR-17, which functions as an oncogenic master regulator targeting 22 genes. We performed extensive functional assays with these master regulatory miRNAs in glioma cell lines and patient-derived stem cells and demonstrated their ability to inhibit cell proliferation, invasion, neurosphere formation, and xenografted tumor growth. Importantly, we also demonstrated that these miRNAs simultaneously targeted multiple glioblastoma-regulating genes in different pathways, decreasing their protein levels. We then used Magnetic Imaging-guided Focused Ultrasound (MRgFUS) in conjunction with microbubbles to transiently open the blood-brain barrier, enabling the local delivery of brain penetrating nanoparticle-conjugated miR-340 into pre-established glioblastoma xenografts. This approach significantly reduced glioblastoma growth and improved mouse survival. This work describes a conceptually new approach to glioblastoma therapy to identify master regulatory miRNAs and use MRI-guided FUS-MB and nanoparticles to deliver these microRNAs in vivo to inhibit glioblastoma growth. Given the availability of clinical grade FUS and BPN for brain applications, these findings pave the way for novel miRNA therapeutic clinical trials.

## Results

### Identification of genome-wide miRNA targets in glioblastoma via PAR-CLIP

Several online miRNA target prediction tools are available, but they can yield false positives and do not always capture all relevant targets. To identify master regulatory miRNAs in glioblastoma, we therefore first used Photoactivatable Ribonucleoside-Enhanced Crosslinking and Immunoprecipitation (PAR-CLIP) to experimentally identify genome-wide targets for all miRNAs in glioblastoma cells (**Fig. 1A**). PAR-CLIP was conducted in U87 cells overexpressing either FLAG-tagged AGO1, AGO2, or AGO3. The cells were cultured in presence of 4-thiouridine and the crosslinked RNA fragments bound by AGO1, AGO2, AGO3 were selectively immunoprecipitated with anti-FLAG antibodies. Crosslinked RNAs were end-labeled using P-32, and immunoprecipitates were fractionated by SDS-PAGE. Ribonucleoproteins (RNPs) were visualized by phosphorimaging, and the RNP corresponding to AGO was isolated and crosslinked RNA fragments recovered, reverse transcribed, and deep-sequenced. Sequence reads were mapped to the human genome and grouped into clusters using PARalyzer to identify those enriched in T-to-C mutations, which are induced by crosslinking of 4SU-containing RNA (**Fig. 1B-C, Fig. S1A-E**). Resulting clusters of sequence reads were then analyzed to identify sites compatible with canonical seed pairing of miRNAs. We observed similar binding patterns for all three human AGO proteins (**Fig. 1D,E**). A total of 19,483 clusters were obtained from the three AGO PAR-CLIP libraries, with 6,412 for AGO1, 7,007 for AGO2, and 6,064 for AGO3. 4,068 clusters were common across all three AGO samples (**Fig. 1D, Table S1**). Among them, 279 mapped to 5′ untranslated regions (5′-UTRs), 4,635 to coding sequences (CDSs), 4,750 to 3′ untranslated regions (3′-UTRs), and 513 to intronic regions (**Fig.1E**). These clusters correspond to 4,583 transcripts, representing approximately 23% of all protein-coding genes. A comprehensive list of mRNA targets for AGO1, AGO2, AGO3, along with their associated miRNAs, is provided in **Table S1**. These findings identify all mRNA targets of all miRNAs expressed in glioblastoma cells. They offer valuable insights into the genome-wide interactions between miRNAs and their targets in glioblastoma cells, illuminating the complex regulatory networks through which miRNAs modulate multiple targets.

**Figure 1:**
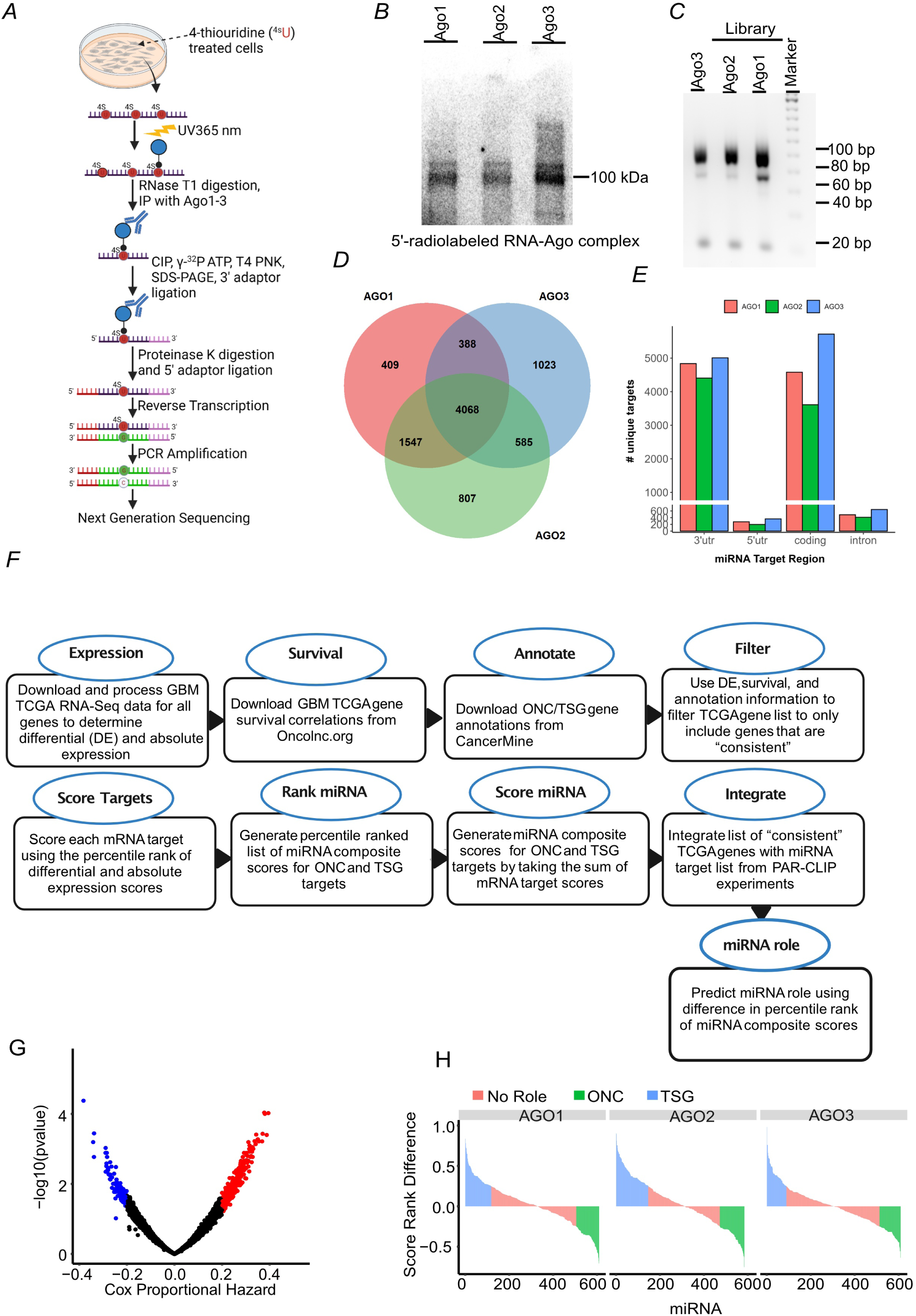
Identification and prioritization of master regulatory miRNAs in glioblastoma: A) Schematic overview of PAR-CLIP. B) The phosphorimage of SDS-PAGE gel of RNA-Argonaute (Ago) complexes labeled with 5′-32P that were immunoprecipitated with a Flag-tag antibody. The complex is expected to appear near 100 kDa. C) Agarose gel separation of PCR products from AGO1, AGO2, AGO3 PAR-CLIP cDNA libraries. D) A Venn diagram illustrating the overlap in miRNA targets identified with AGO1, AGO2, AGO3. E) Bar graph showing the distribution of identified targets within coding and 3’-UTR regions. F) Diagram depicting the algorithm used for determining master regulatory miRNAs in glioblastoma. G) Volcano plot denoting the significant miRNAs based on Cox Propportional Hazard. H) Classification of miRNAs as oncogenic and tumor-suppressive based on their targets derived from AGO1, AGO2, AGO3 PAR-CLIP.

### Ranking of master regulatory miRNAs based on the importance of their targets in human TCGA tumors

Having identified all targets of all miRNAs in glioblastoma cells, we next sought to determine the relevance of these targets, and consequently of their targeting miRNAs, to glioblastoma malignancy. This process is described in the methods section (**Fig. 1F**). Briefly, we analyzed The Cancer Genome Atlas (TCGA) data for the PAR-CLIP-identified miRNA targets in human tumors. We identified significantly deregulated targets (FDR ≤ 0.05) with a greater than 2-fold change in human tumors using the TCGA database. The targets were then scored based on the magnitude and frequency of dysregulation, as well as their correlation with patient survival, using a Cox coefficient threshold of 0.3 (**Fig. 1G, Table S2 & 3**). Each target gene was assigned a composite score calculated from its absolute expression level in glioma samples and its fold change relative to normal brain tissue. These two metrics were independently converted into percentile ranks (ranging from 0 to 1), and their sum represented the gene’s composite score, with a theoretical range from 0 (lowest expression and change) to 2 (highest in both categories). For each miRNA, two aggregate scores were then computed: a tumor suppressor gene (TSG) score, representing the average composite score of its oncogenic targets, and an oncogene (ONC) score, reflecting the average composite score of its tumor suppressive targets. To enable comparison across the entire miRNA dataset, these scores were further normalized by converting them into percentile ranks across all miRNAs. To determine the functional role of each miRNA, we calculated the difference between its TSG and ONC percentile ranks. We then added the scores for each target, and then added all scores for all targets of each miRNA. We subsequently averaged the target scores for both tumor suppressor and oncogenic miRNAs across the AGO1, AGO2, and AGO3 PAR-CLIP datasets, and consolidated the comprehensive set of targets for each miRNA (**Table S4 & 5**). The final score for each miRNA represents is potential master regulatory and therapeutic potential because it is a reflection of both the number of its targets and the importance of these target in glioblastoma biology.

Using this computational workflow, we identified several miRNAs that targeted many highly relevant genes that are dysregulated in GBM (**Fig.1H, Fig. S1F** and **Table S1-5)**. We designated these miRNAs as master regulators because they can simultaneously target multiple significantly deregulated and biologically relevant genes and inhibit their expressions. This new approach identifies miRNAs that are likely to exert strong therapeutic effects when delivered (tumor suppressive miRNAs targeting numerous oncogenes) or inhibited (oncogenic miRNAs that target numerous tumor suppressors). The comprehensive list of miRNAs, their targets, and their scores can be found in **Table S1-5**.

### Master regulatory miRNAs miR-340 and miR-382 bind multiple targets and decrease their expression

To prioritize master regulatory miRNAs for functional validation, we incorporated evolutionary conservation as an additional criterion and selected the top five master regulatory miRNAs from both the tumor suppressor (Table S4) and oncogenic (Table S5) lists for further analysis. Among these, we focused on two tumor-suppressive miRNAs, miR-340 and miR-382 because these miRNAs had high scores based on the algorithm described above. Our analysis determined that the tumor-suppressive master regulatory miRNAs, miR-340 and miR-382, have 91 and 79 targets (**Table S4**), respectively. From these, we selected a subset for validation based on their known deregulation in glioblastoma and other cancers and their widespread expression in glioblastoma samples and patient-derived glioblastoma stem cells. We checked the expression of these selective targets, CD44, TOP2A, MDM2, RHOC, HMGA2, PLAU, and NUSAP1 in TCGA glioma patient samples and found that they are highly overexpressed in tumor samples compared with GTEx normal samples (**Fig. S2A-G**). We performed extensive target validation using immunoblot and 3’UTR reporter analyses. We transfected precursor miR-340 and miR-382 into glioma cell lines A172, U87, U251, as well as patient-derived glioma stem cell lines GSC-34 and GSC-28. Cell lysates were then subjected to immunoblot. Overexpression of these miRNAs in various glioblastoma cells, and stem cells, led to a marked reduction in protein levels across all tested cell lines (**Fig. 2A-I**). To determine if miR-340 and miR-382 directly target CD44, TOP2A, RHOC, MDM2, HMGA2, EGFR, PDGFRA, NUSAP1, and PLAU by binding to their 3’ untranslated regions (3’UTRs), we amplified the 3’UTR sequences containing the miR-340 and miR-382 binding sites from genomic DNA and cloned them into the psiCheck2 luciferase reporter plasmid under the control of the T7 promoter. These 3’UTR luciferase reporter constructs were then transiently transfected into U87 or 293T cells along with either a scrambled control miRNA mimic or mimics of miR-340 or miR-382. Overexpression of miR-340 resulted in a significant 3-fold to 5-fold decrease in luciferase signals for *CD44, TOP2A, RHOC, HMGA2, MDM2, EGFR* and *PDGFRA* (p < 0.05). Similarly, miR-382 overexpression led to a significant 3-fold to 5-fold reduction in 3’UTR luciferase signals for PLAU, CD44, NUSAP1, HMGA2, and MDM2 (p < 0.05), compared to scrambled control transfections (**Fig. 2J,K**). These findings provide strong evidence that miR-340 and miR-382 directly bind to their respective 3’UTRs, significantly repressing target gene expression.

**Figure 2:**
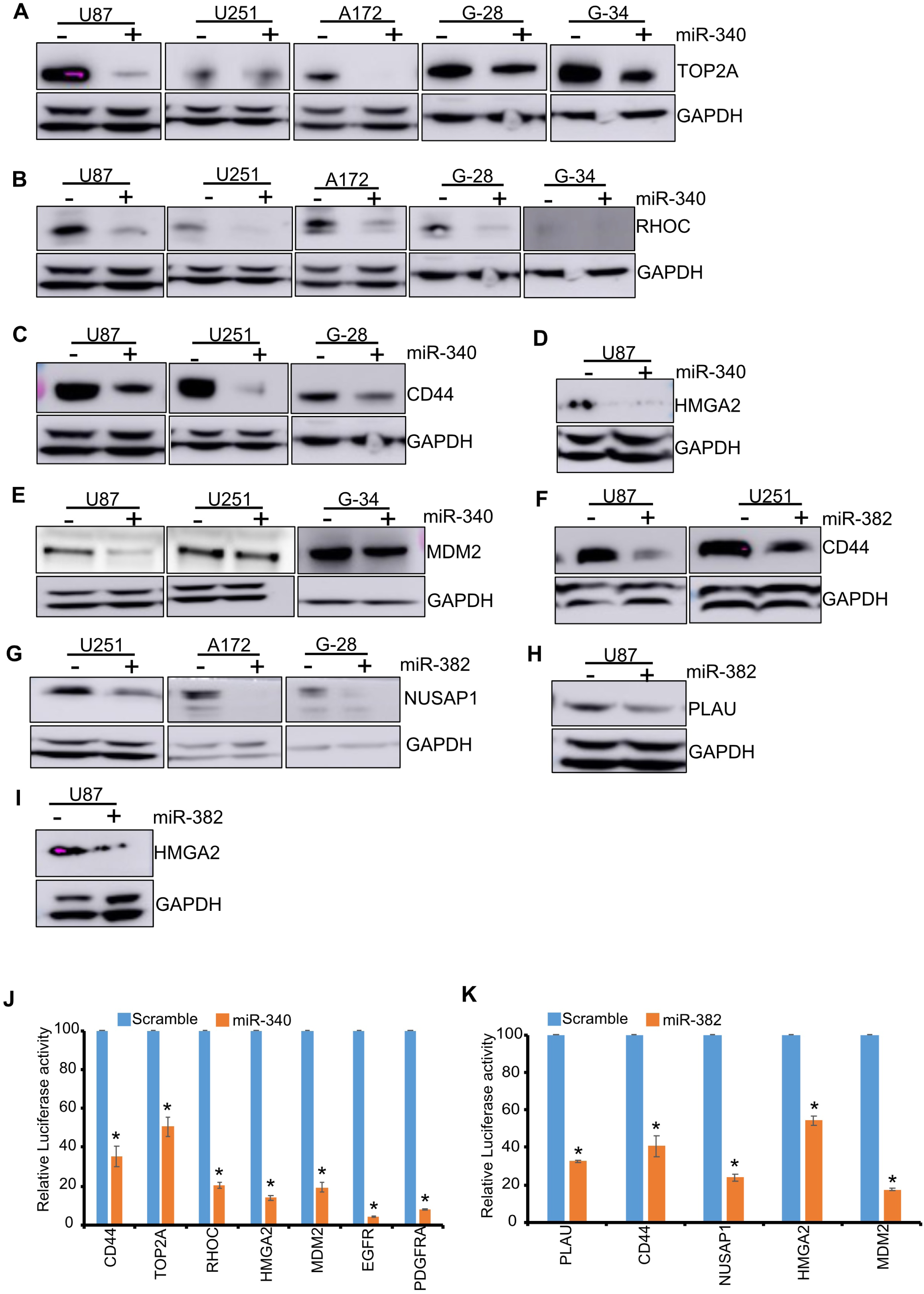
Validation of miR-340 and miR-382 targets in glioblastoma cells and patient-derived stem cells: A) Glioblastoma cell lines A172, U87, U251, and patient-derived stem cell lines GSC-28 and GSC-34 were transfected with either a scrambled negative control, miR-340, or miR-382. Immunoblots were probed with antibodies against TOP2A (A), RHOC (B), CD44 (C), HMGA2 (D), and MDM2 (E), CD44 (F), NUSAP1 (G), PLAU (H), and HMGA2 (I). GAPDH served as the internal loading control. The data show that miR-340 and miR-382 downregulated protein expression compared to negative controls. J,K) 3’UTR Luciferase activity assay in U87 cells were performed by co-transfecting cells with miR-340 (J) or miR-382 (K) and a psiCheck2 luciferase reporter plasmid containing the 3’-UTR regions of targets CD44, TOP2A, RHOC, HMGA2, MDM2, EGFR, PDGFRA (J), or PLAU, CD44, NUSAP1, HMGA2, and MDM2 (K). The data show that miR-340 and miR-382 decreased luciferase activity for all respective targets compared to controls. Statistical significance was determined using a two-tailed Student’s t-test, with * = P<0.05.

### miR-17 is an oncogenic master regulatory miRNA in glioma

We identified miR-17 as one of the top oncogenic master regulatory miRNAs that targets several deregulated tumor suppressor genes in glioblastoma. We determined the expression of the selective targets in TCGA glioma samples and found they are highly downregulated in tumor samples compared with GTEx normal samples (**Fig. S3A-D**). We validated some of these targets using immunoblot analysis and 3’UTR reporter assays and in several glioma cell lines and patient-derived glioma stem cell lines. Since miR-17 is upregulated in glioblastoma and acts as an oncogene, we inhibited miR-17 by transfecting an anti-miR-17 inhibitor into glioma cell lines (A172, U87, U251) and patient-derived glioma stem cell lines (GSC-28 and GSC-34). We selected specific targets— *ZBTB4, ANKRD11, EHD3,* and *EPHA4* based on their widespread published roles in glioblastoma. Inhibiting miR-17 resulted in an increased expression of all these targets in both glioblastoma cells and stem cells, indicating that miR-17 suppresses their expression (**Fig. 3A-D**). To confirm that miR-17 directly binds to the 3’UTRs of *ANKRD11, EHD3, EPHA4,* and *ZBTB4,* we conducted a luciferase reporter assay. The 3’UTR regions of these genes, which contain the miR-17 binding sites, were cloned into a *psiCheck2* reporter plasmid. Glioblastoma cells were transfected with either a scrambled anti-miR or an anti-miR-17 inhibitor, along with the luciferase reporter plasmid containing the miR-17 target binding sites. Transfection with the miR-17 inhibitor led to a significant de-repression (increase) in luciferase signals for *ANKRD11, EHD3, EPHA4,* and *ZBTB4* (p < 0.05), compared to the scrambled control (**Fig. 3E**). These data show that miR-17 directly binds to and regulates several deregulated mRNAs in glioblastoma.

**Figure 3:**
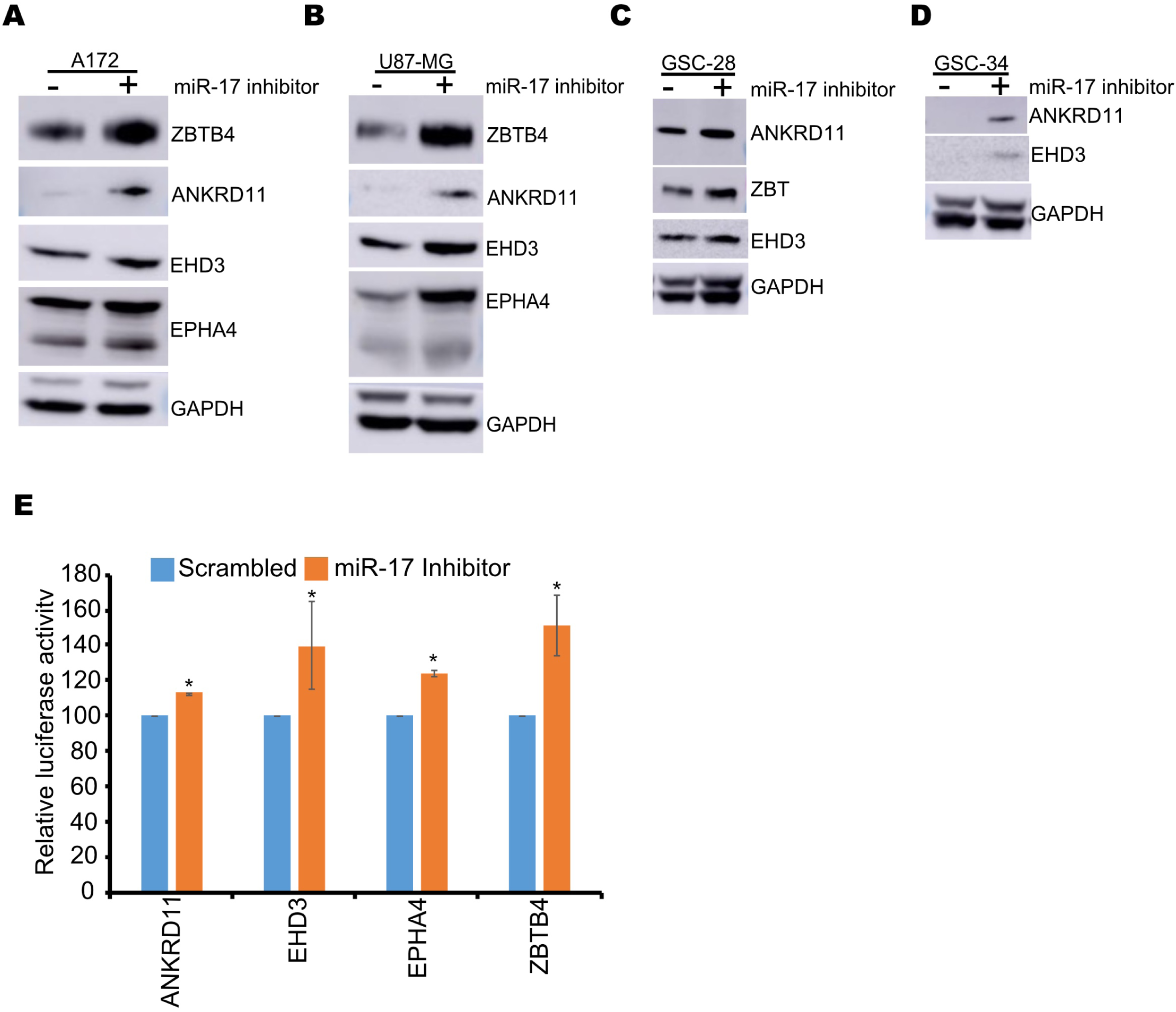
Validation of miR-17 targets in glioblastoma cells and patient-derived stem cells: A-D) Glioma cell lines A172, U251, GSC-28, and GSC-34 were transfected with a miRNA inhibitor targeting miR-17. Immunoblots were performed with antibodies against ZBTB4, ANKRD11, EHD3, and EPHA4. GAPDH served as the internal control for loading. The data show that the miR-17 inhibitor increased protein expression compared to negative controls. E,F) 293T cells were co-transfected with miR-17 or anti-miR-17 along with psiCheck2 luciferase reporter plasmid containing the 3’-UTR regions of targets ANKRD11, EHD3, EPHA4 and ZBTB4. The data show that the miR-17 inhibitor increased luciferase activity for all respective 3’UTR targets compared to negative controls 48 hours after the transfection, cells were lysed and luciferase signals were measured. * = P<0.05.

Further, we investigated whether the targets of the master regulatory microRNAs, miR-340, miR-382, and miR-17, are associated with patient survival. To evaluate survival outcomes, we utilized two independent cohorts of primary glioma patient samples from The Cancer Genome Atlas (TCGA) and the Chinese Glioma Genome Atlas (CGGA) for survival analysis. Kaplan-Meier survival curves were generated and analyzed using the online tools GEPIA and CGGA, where patients were stratified into high and low gene expression groups based on target expression levels.For the tumor suppressor targets of miR-340 and miR-382, high expression was correlated with poorer patient survival, while low expression was associated with more favorable outcomes across both TCGA and CGGA cohorts (Fig S4A-G & Fig S5A-G). Conversely, for the oncogenic miR-17, high expression of its targets was linked to better survival, while low expression was associated with poorer patient prognosis (FigS6A,B & FigS7A,B)Additionally, we performed survival analysis using CGGA data from recurrent glioma patient samples, which revealed similar survival trends for both tumor suppressor and oncogenic miRNA targets (FigS8A-F & FigS9). These findings underscore the significance of master regulatory miRNAs and their target genes in determining glioma patient outcomes.

### Master regulatory miRNAs miR-340 and miR-382 regulate multiple pathways in glioblastoma

After identifying and validating master regulatory miRNAs and their targets, we next investigated their functions. We first performed pathway analyses with the targets of miR-340 and miR-382. We performed gene ontology (GO) term, Kyoto Encyclopedia of Genes and Genomes (KEGG), and hallmark pathway analysis. The most significant pathways associated with miR-340 were the mitotic cell cycle phase transition, stem cell differentiation, miRNAs in cancer, PI3K-Akt signaling, proteoglycan in cancer, the hallmarks of apoptosis, G2/M-checkpoint and epithelial to mesenchymal transition (**Fig. 4A-C**). The pathways enriched for miR-382 targets were response to hypoxia, signaling pathways associated with miRNA metabolic process, proteoglycan in cancer, miRNAs in cancer, G2/M checkpoint, and epithelial to mesenchymal transition (**Fig.4D-F**). These data confirm that these miRNAs regulate different cancer-related pathways and inhibiting them could attenuate glioma growth by inhibiting numerous oncogenic pathways.

**Figure 4:**
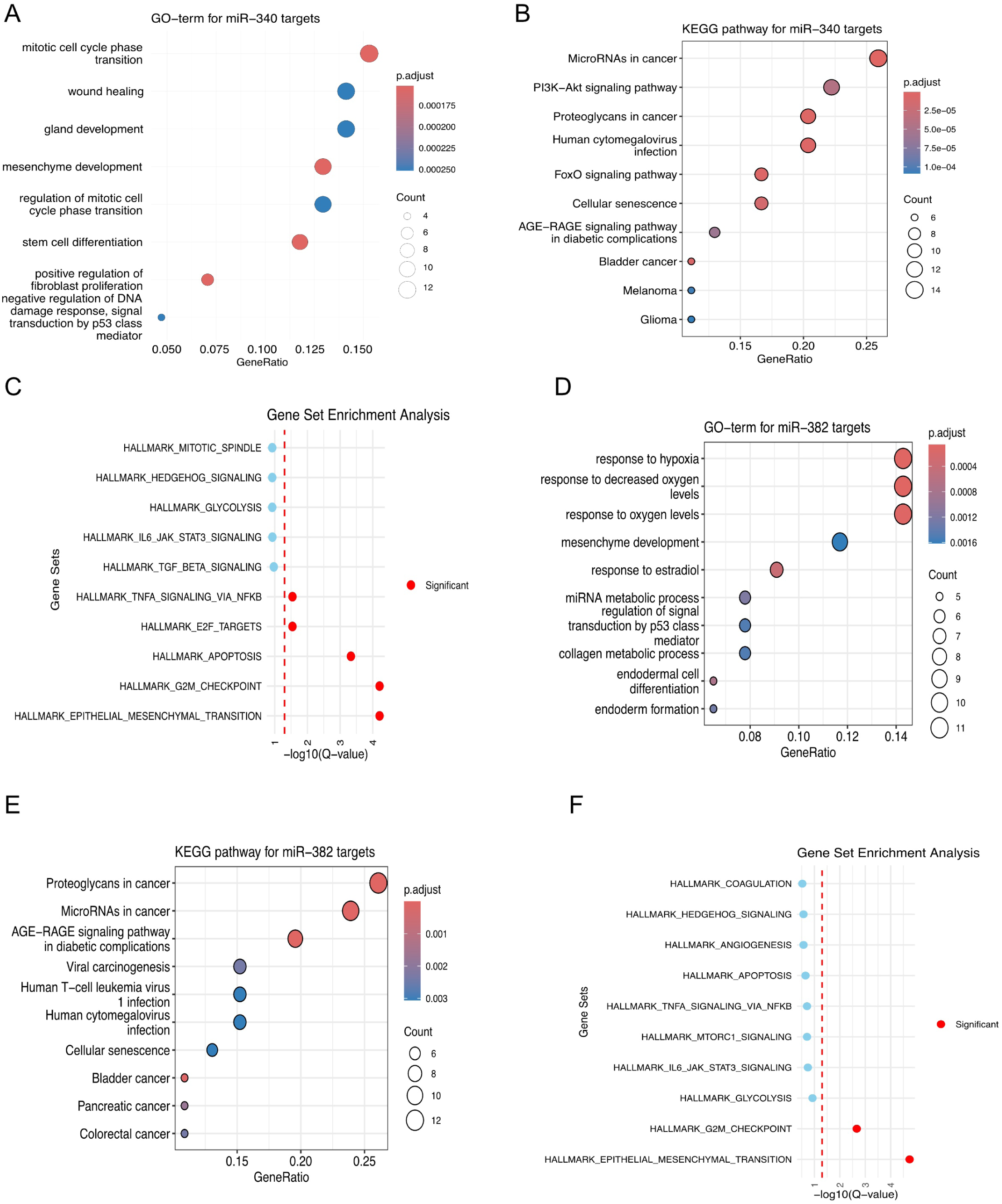
miR-340 and miR-382 regulate cancer pathways: A-F) Targets identified for miR-340 and miR-382 through PAR-CLIP and The Cancer Genome Atlas (TCGA), were utilized as input for pathway analysis involving Gene Ontology (GO) terms for Biological Processes (BP), Kyoto Encyclopedia of Genes and Genomes (KEGG), and gene set enrichment analysis. The significant pathways are highlighted by red dots.

### Master regulatory miRNAs miR-340 and miR-382 inhibit cell proliferation, invasion, and neurosphere formation in several glioblastoma cell lines

We then performed a thorough assessment of the functional and experimental therapeutic impacts of miR-340 and miR-382. We first quantified their endogenous expression levels in glioblastoma cell lines, GSCs, and banked human tumors including cell lines (A172, T98G, LN18, U251, U87, SNB19), and stem cell lines (GSC-28, GSC-20, GSC-34, GSC-627, GSC-267), as well as patient glioblastoma samples, using quantitative PCR. Normal human astrocytes and normal human cortex were used as a controls. We observed a significant downregulation of miR-340 and miR-382 expression across all cell lines compared to normal human astrocytes, mirroring the pattern observed in patient glioblastoma samples compared to normal brain samples. This consistent downregulation hinted at a potential tumor suppressor role for these miRNAs, which was consistent with our analyses (**Fig. S10A-D**).

Subsequently, we conducted cell proliferation and invasion assays to determine the biological significance of miR-340 and miR-382 *in vitro* across different glioblastoma cell lines. Glioblastoma cells were transfected with either scrambled control miRNA or precursor miR-340 or precursor miR-382, and cell proliferation was assessed through trypan blue staining and live cell counting. Overexpression of precursor miR-340 or miR-382 led to a significant reduction in cell proliferation compared to the scrambled negative control (**Fig. 5A-D**). To investigate the influence of these miRNAs on glioma cell invasion, we performed transwell invasion assays using glioblastoma cell lines. The transwell invasion chambers were pre-coated with collagen IV, one of the abundant extracellular matrix components in the brain. Glioblastoma cell lines were transfected with scrambled control miRNA, or miR-340, or miR-382 precursors before being plated in the transwell chambers. These cells were allowed to invade through the collagen IV layer. Transient overexpression of either miR-340 or miR-382 led to a substantial reduction in glioblastoma cell invasion compared to scrambled negative control miRNA-transfected cells (**Fig.5E-J**).

**Figure 5:**
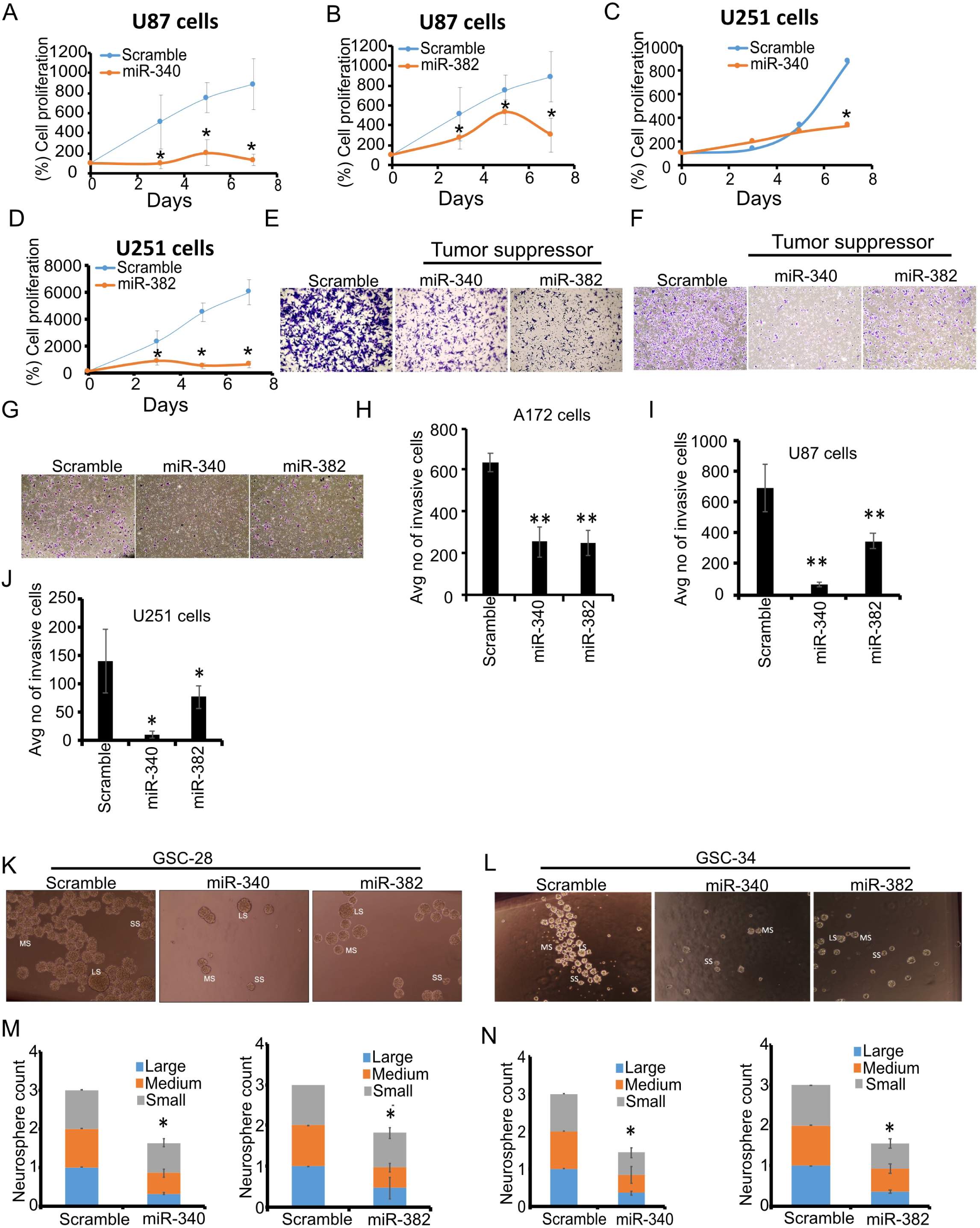
miR-340 and miR-382 inhibit cell proliferation, invasion, and neurosphere formation in glioblastoma: A-D) Glioblastoma cell lines U87 and U251 were transfected with either a scrambled control, or miR-340, or miR-382. Cell counts were performed at various time points, and miR-340 and miR-382 showed decreased cell growth compared to negative controls. E-J) A172, U87, and U251 cells were transfected with mimic miR-340 and miR-382 and invasion assays were executed. Invaded Images from 5-10 random fields were captured and analyzed using ImageJ software for quantification. miR-340 and miR-382 decreased invasion compared to negative controls K-N) Six-well plates were pre-coated with poly-ornithine, and glioblastoma stem cell lines GSC-28 and GSC-34 were plated and transfected with either scrambled control miRNA, miR-340, or miR-382. Neurosphere images were taken from five distinct fields 72 hours post-transfection and categorized into large, medium, and small using ImageJ software, with quantifications presented in (M, N). miR-340 and miR-382 decreased neurosphere formation compared to negative controls. Data represent mean ± SEM from three independent experiments. * = P<0.05.

Glioblastoma contains self-renewing stem cells contributing to tumor initiation and resistance to therapy (*45*). To elucidate the effect of the master regulatory miRNAs on the self-renewal property of glioblastoma stem cells, we conducted neurosphere assays using two distinct patient-derived glioblastoma stem cell lines, GSC-34 and GSC-28. We transfected precursor miR-340 or precursor miR-382 or scrambled controls into the glioblastoma stem cells and counted the number of neurospheres seven days post-miRNA-transfection. Neurospheres were categorized based on their size under the microscope (10 X magnification) into large, medium, and small groups. Overexpression of miR-340 and miR-382 in these glioma stem cells led to a reduction in the overall number of neurospheres, indicating that these master regulatory miRNAs diminish the self-renewal capabilities of glioblastoma stem cells (**Fig. 5K-N**).

### Inhibiting oncogenic miR-17 in glioma cells decreased cell proliferation, invasion, and neurosphere formation in a spectrum of glioma cell lines

miR-17 is one of the top oncogenic master regulatory miRNAs identified by our integrated approach. Similar to the tumor suppressive master regulatory miRNAs, we first checked the endogenous expression of miR-17 in multiple glioma and stem cell lines by RT-qPCR **(Fig. S10E,F)**. We observed an increased expression pattern in all glioma and stem cell lines consistent with our algorithm that identified miR-17 as an oncogenic miRNA. Therefore, we performed a series of functional assays with miR-17 in a spectrum of glioblastoma cell lines, including patient-derived glioblastoma stem cells. For the cell proliferation assay, we transfected the inhibitor against miR-17 into multiple glioma cell lines, and counted the cells on different days. We observed a decrease in cell proliferation supporting the oncogenic nature of miR-17 in glioma cells **(Fig. 6A-C)**.Transwell cell invasion assays were carried out with anti-miR-17 in glioblastoma cell lines. The invasion assays wwere carried out with a collagen IV coated transwell chamber. Inhibition of miR-17 in glioblastoma cell lines led to a decrease in cell invasion through collagen-coated chambers, which were counted by taking images from five random microscopic fields and quantified with ImageJ software (**Fig. 6D-I)**. The neurosphere formation assay was performed with glioma stem cell lines GSC-28 and GSC-34. The transfection of glioma stem cell lines with an inhibitor of miR-17 decreases the overall neurosphere formation. We categorized the neurosphere size into three different groups, large, medium, and small, based on the size calculator with ImageJ software. The overall neurosphere number in different categories decreased significantly after miR-17 inhibitor treatment compared to the scrambled control treatment **(Fig. 6J-M)**.

**Figure 6:**
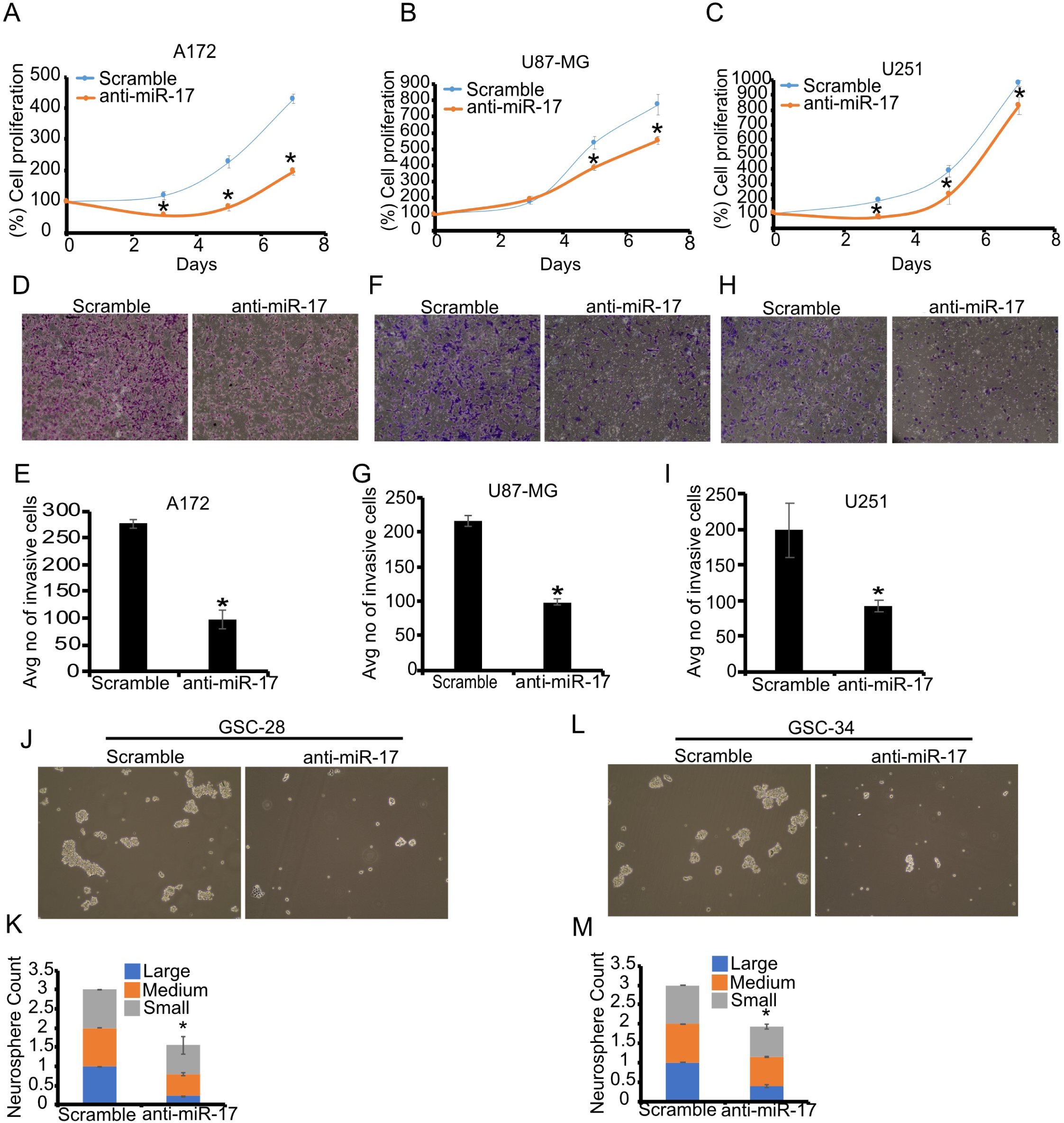
Effects of miR-17 inhibition on cell proliferation, invasion, and neurosphere formation in glioblastoma: A-C) A172, U87, and U251 were transfected with either scrambled control miRNA or a miR-17 inhibitor. Cells were counted 48 hours post-transfection using trypan blue exclusion at various intervals to assess viability. D-I) A172, U87, and U251 cells were transfected with control or miR-17 inhibitor. Invasion assay was carried out and the invaded cells were stained with crystal violet, and images were captured and analyzed using ImageJ software for quantification. J-M) Glioma stem cell lines GSC-28 and GSC-34, plated on poly-ornithine-coated 6-well plates, were transfected with either a scrambled control miRNA or an miR-17 inhibitor. Neurospheres were imaged 72 hours post-transfection in the neurobasal complete growth medium. Images from five different microscopic fields were taken, and neurosphere sizes were categorized into large, medium, and small for quantification using ImageJ software. Data are presented as mean ± SEM from three independent experiments. * = P<0.05.

### miR-340 and miR-382 inhibit *in vivo* glioblastoma growth

The impact of miR-340 and miR-382 overexpression on xenografted tumor growth in mice was investigated.. We transfected 3×10^5^ U87 cells with either a scrambled negative control miRNA (n=7), precursor miR-340 (n=7), precursor miR-382 (n=7), or anti-miR-17 (n=7) **Figure 7A**. The transfected cells were then stereotactically implanted into the striata of 6-week-old immunodeficient mice. Over three weeks, the mice were closely monitored for tumor growth and survival, and MRI images were obtained to visualize the tumors.

**Figure 7:**
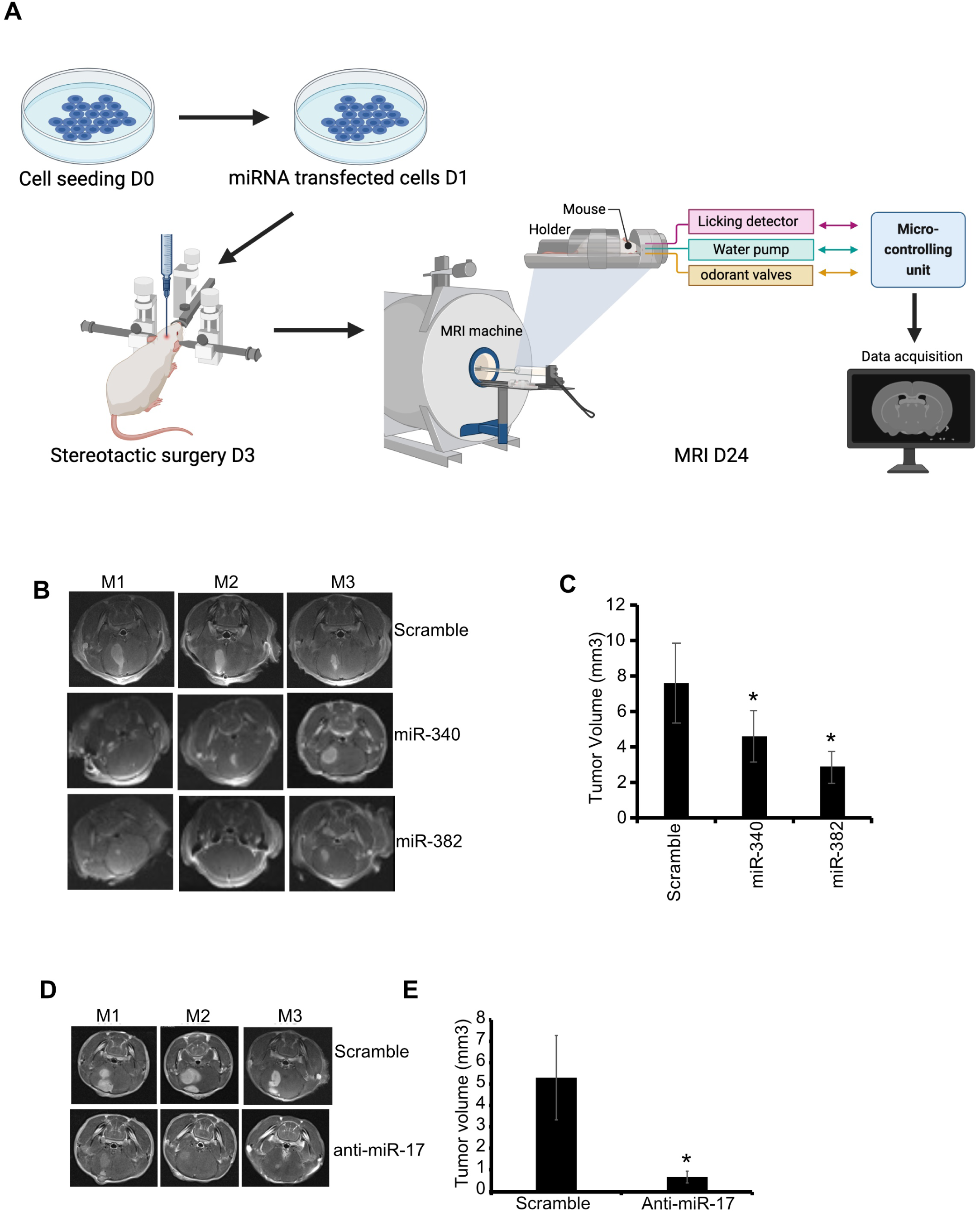
miR-340, miR-382 and miR-17 regulate in vivo tumor growth: A) Schematic overview of the experimental design for tumor implantation and timeline for MRI imaging to assess tumor volume. Each group comprised seven mice for surgery. B-E) U87 cells were transfected with scrambled negative miRNA, miR-340, miR-382, or an inhibitor of miR-17. 48 hours post-transfection, cells were implanted intracranially into the striata of 5-6-week-old immunodeficient mice. Mice were monitored for 3-4 weeks, and MRI imaging was performed to evaluate tumor generation. Representative MRIs and quantification showing that miR-340, miR-382 and anti-miR-17 inhibitor showed reduction in tumor volume compared to Scramble controls. * = P<0.05.

After three weeks, the scrambled control U87 group exhibited significant tumor growth. In contrast, the groups treated with miR-340 and miR-382 displayed a significant reduction in tumor volume **(Fig. 7B, C)**. Specifically, the scrambled control group reached a tumor volume of (7.58 ± 2.26) mm^3^, while the mice bearing miR-340- or miR-382-transfected tumors showed tumor volumes of (4.59 ± 1.44) mm^3^ and (2.86 ± 0.9) mm^3^, respectively (**Fig. 7C**). The mice implanted with miR-17 inhibitor transfected cells showed a substantial decrease in tumor volume compared to scrambled control **(Fig. 7D,E)**. The scrambled control mice group showed tumor volumes of (5.3 ± 1.94) mm^3^, whereas miR-17 inhibitor group exhibited tumor volumes of 0.68 +0.29 mm^3^. Collectively, these *in vivo* findings support the use of miR-340, miR-382, the miR-17 inhibitor, or a combination of them as novel therapeutics for glioblastoma.

### Therapeutic delivery of nanoparticle-conjugated miR-340 to mice bearing glioblastoma tumors by MRI-guided Focused Ultrasound inhibits tumor growth

For the therapeutic delivery of master regulatory miRNA, we developed and employed a novel approach consisting of MRI-guided focused ultrasound (FUS) and microbubbles (MB) (FUS-MB) to facilitate the delivery of lentiviral vectors encoding for miR-340 that were conjugated with brain-penetrating nanoparticles (BPN). This innovative approach (abbreviated FUS-MB-BPN) circumvents the hurdles of miRNA therapeutics, particularly the blood-brain barrier and tumor penetration and transfection, and delivers cargo loads to tumor cells. The schematic plan for FUS-MB-BPN is depicted in **Figure 8A-B**. We generated glioblastoma xenografts in immunodeficient mice, accomplished by intracranial implantation of 3×10^5^ U87 cells into the striata of mice brains. Once the tumors had formed (7-10 days post-injection), we verified tumor formation by MRI and the safe and effective opening of the blood-brain barrier. This step was achieved through a meticulously orchestrated combination of microbubbles and sonication, resulting in a conspicuous increase in MRI signal intensity surrounding the tumors. This increased signal intensity is attributed to the leakage of the MRI contrast agent, gadobenate dimeglumine, into the brain parenchyma after blood-brain barrier opening (**Fig. 8C**). To deliver miR-340 into brain tumors, we synthesized poly (beta-amino esters) (PBAE) nanoparticles and coated their surface with polyethelene glycol (PEG). The PEGylated nanoparticles were mixed with CMV promoter driven miR-340 lentiviral plasmid DNA. We also prepared the BPN particles with control plasmid encoding a scrambled sequence. We extensively characterized these BPN particles because the zeta potential and size of the particles are important parameters for effective blood-brain barrier penetration. The zeta potential and average size of the miR-340 BPN were 70.54 nm and 52.15 nm, respectively, whereas for control, they were 72.51 nm and 51.62 nm, respectively. The mir-340 or control BPN were intravenously injected alongside microbubbles (MB), and focused ultrasound (FUS) was applied to the tumor regions under MRI guidance to transiently open the blood-brain barrier and facilitate the delivery of the -BPN particles into the tumor. We divided our brain tumor-bearing mice into two cohorts: one receiving nanoparticle conjugated scrambled control miRNA and the other treated with the tumor-suppressive miR-340. Each group consisted of seven mice. One week after the FUS-MB-BPN procedure, MRI images were taken to visualize and measure the tumor volumes. Delivery of miR-340 by FUS-MB-BPN into the tumors significantly reduced tumor burden compared to the control group (n = 7 mice per group, *=P<0.05). This represents the first successful BPN-FUS-based experimental therapeutic delivery of a miRNA to inhibit tumor growth (**Fig. 8D-E**). We also closely monitored the post-FUS-treated mice for survival, which revealed miR-340 delivery into brain tumors significantly extended the survival of these mice compared to those treated with the scrambled control miRNA (**Fig. 8F**). To assess potential toxicity from FUS-MB, we harvested various organs, including the liver, kidney, brain, heart, spleen, lung, lymph nodes and pancreas from treated mice. We then performed H&E staining on these organs, which were analyzed by a Neuropathologist. The data showed no apparent damage to the liver, kidney, brain, or heart in treated mice compared to the non-treated controls (**Fig. S11A,B**). These data demonstrate that MRI-guided FUS-MB-BPN is a safe and effective strategy for delivering miRNA into brain tumors.

**Figure 8:**
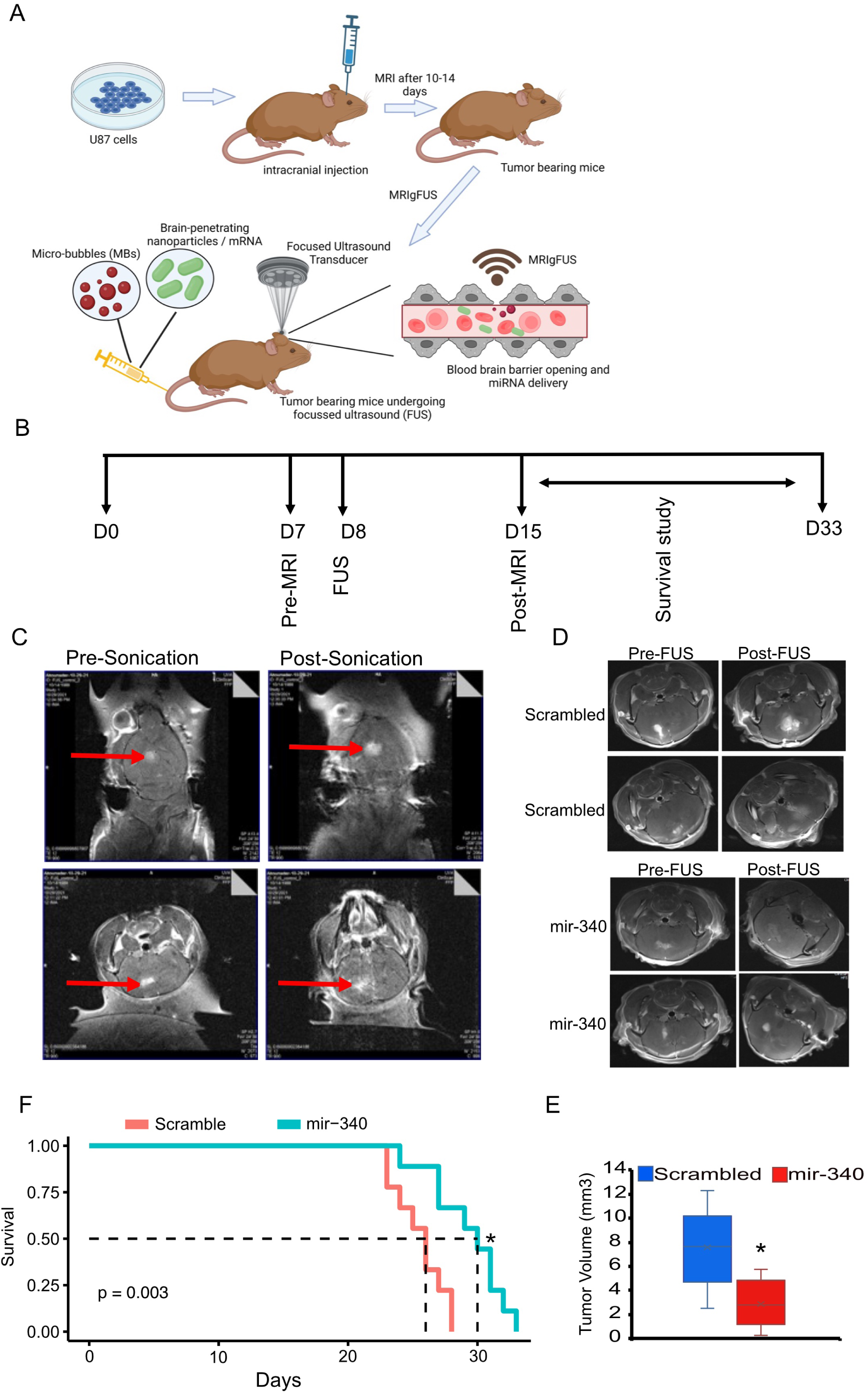
Inhibition of in vivo glioma growth through MRI-guided (MRIg) Focused Ultrasound, Microbubbles and Brain-Penetrating Nanoparticles (FUS-MB-BPN) delivery of miR-340: A) A schematic representation of FUS-MB-BPN-mediated miRNA delivery into mice. B) Overall experimental plan and timeline for FUS-MB-BPN. C) Tumors in mice were sonicated pre and post FUS-MB-BPN, and images were captured to validate blood-brain barrier opening. MB were employed to facilitate blood-brain barrier opening. Arrows indicated the tumor location and show dispersion of the contrast demonstrating successful opening of the blood-brain barrier. D,E) Microbubbles were injected through the mice’s tail vein, and MRIgFUS was conducted. Upon confirmation of blood-brain barrier opening, BPN conjugated with either scrambled or miR-340 were injected through the mice’s tail vein. The mice were imaged using MRI, and tumor volume shows significant reduction upon delivery of miR-340 compared to Scramble controls at day 15. F) Kaplan Meir survival curve showing miR-340 significantly prolonged survival compared to scrambled control mice. n = 7 mice per group. *=*P* < 0.05 for miR-340-treated vs. scramble control.

## Discussion

Glioblastoma is a lethal, and aggressive primary brain tumor (*46*). Despite extensive research and clinical trials, targeted therapies have faced insurmountable challenges presented by the blood-brain barrier, tumor invasiveness, resistance to cytotoxic therapies and tumor heterogeneity (*47*) (*48*). One major hurdle towards the success of targeted therapies for glioblastoma is the concurrent deregulation of numerous genes in any single tumor (14, 15). The simultaneous targeting of several deregulated oncogenic drivers using conventional drugs is severely limited by the fact that the drugs needed to simultaneously target many molecules do not currently exist, and because combining several drugs in a clinical setting leads to an exponential increase in toxicity (20). To overcome these limitations, we developed and successfully tested a new miRNA therapeutic strategy.

MicroRNAs are small noncoding RNAs that can be deregulated in cancers and brain tumors, where they can function as either oncogenes or tumor suppressors (*49, 50*). Because miRNA do not require full complementary with the targeted mRNA sequences to inhibit gene expression, a single miRNA can target multiple genes and inhibit their expression (*51*). We therefore reasoned that there exist master regulatory miRNAs, which can inhibit multiple deregulated genes in glioblastoma, and that these miRNAs can be used as therapeutic agents/targets, equivalent to using multiple drugs in combination (*52–54*). Combination drug therapies are difficult because many deregulated molecules do not have drugs that target them, and because combining several drugs can lead to exponential increase in toxicity (*55, 56*). To find master regulatory miRNAs, we developed an approach that integrates miRNA target identification with PAR-CLIP with TCGA data analyses to find miRNAs that target multiple dysregulated important genes in glioblastoma. This unique approach was invented and developed in our lab and has not been described or published before. It can be applied to any human cancer.

Our approach uncovered numerous miRNAs that had numerous deregulated targets in glioblastoma. We focused on the highly ranked tumor suppressive miR-340 and miR-382 and the oncogenic miR-17. We validated a few of their targets, TOP2A, RHOC, CD44, HMGA2, MDM2 for miR-340 (**Fig. 2A-E**) and CD44, NUSAP1, PLAU, and HMGA2 for miR-382 targets **(Fig. 2F-I)**, and ZBTB4, ANKRD11, EHD3, and EPHA4 for miR-17. The targets we chose because of their known oncogenic/tumor suppressive roles in multiple cancers (*57–64*). These selected targets were validated across multiple cell lines, including patient-derived glioma stem cell lines, using immunoblotting and 3’UTR reporter assays. CD44 is a cell surface adhesion protein highly expressed in many cancers, including cancer stem cells, and regulates cancer progression and metastasis (*65*). TOP2A has been identified as an oncogene in multiple cancers (*58, 66*). TOP2A is overexpressed in pan-cancers, and overexpression is correlated with poor prognosis and advanced pathological stages in most cancers (*66*). RhoC is a member of the RhoGTPase family protein, which has been shown to be involved in cancer cell migration, invasion, and metastasis (*67*). All selected targets are deregulated in glioblastoma, where they play important roles in regulating malignancy. More targets were identified and validated but are not all discussed in this section due to space limitations.

We then functionally validated the master regulatory miRNAs by performing growth, death, invasion, and differentiation assays as well as in vivo tumor growth assays. These assays validated the tumor-suppressive (miR-340 and miR-382) or oncogenic (miR-17 potential) effects of the selected master regulatory miRNAs. We showed that the impact of these miRNAs was pervasive and profound as illustrated in Figure 4A-F. Given their ability to orchestrate concerted regulation of multiple pathways, we aptly designated miR-340, miR-382, and miR-17 as “master regulators,” and potential therapeutic agents or targets for glioblastoma therapy.

To translate these findings into therapy, we designed and tested a novel approach for the therapeutic delivery of miRNAs to glioblastoma animal models. We conjugated miR-340 plasmid DNA with brain penetrating nanoparticles and employed MRI-guided focused ultrasound in conjunction with microbubbles to deliver this therapeutic payload to the brains of mice harboring glioblastoma. This approach led to a significant reduction in tumor growth and prolongation of animal survival. The mice did not exhibit any signs of toxicity. This new noninvasive delivery method represents a pivotal step towards effective and targeted miRNA therapeutics. Efficient drug delivery into brain tumors is impeded by the blood-brain barrier (BBB). Utilizing MRI-guided Focused Ultrasound (MRIgFUS) combined with microbubbles provides a noninvasive technique to temporarily open the BBB for delivering therapeutic molecules with brain penetrating nanoparticles at the tumor site, offering both temporal and spatial control (*68–70*). MRgFUS combined with microbubbles has been utilized in multiple studies to deliver therapeutic molecules into brain tumor (*69, 71*). The blood brain barrier inhibits majority of chemotherapeutic drugs delivery into the brain including the chemotherapeutic drug doxorubicin (*72*). The polymeric nanoparticles have many advantages over cationic polymers such as polyethylenamine (PEI), poly-L-lysine (PLL) nanoparticles because they have more stability in the bloodstream, superior blood-brain barrier crossing potential, increased drug solubility and more drug encapsulated load and controlled drug release make them more suitable for delivering therapeutics when combined with MFgFUS. Among the promising polymers, poly(β-amino esters) (PBABEs) offer a library of nontoxic, biodegradable materials for the compaction of nucleic acids(*73*). In our study we used surface modified PEGylated PBAE nanoparticles which can penetrate brain parenchyma and open BBB efficiently when combined with microbubbles and FUS (*73, 74*). This blood brain penetrating nanoparticles are more diffusive in nature and were able to circulate in the brain at least for 24 hours suggesting they have low adhesive interactions with extracellular matrix that could lead to the clearance or low distribution of this particles in the brain parenchyma. The precision of focused ultrasound transducers enables them to focus on a millimeter scale, accurately targeting only the tumor region.

In conclusion, our study developed and successfully tested novel approaches to identify and therapeutically deliver miRNAs to glioblastoma. This can be translated into clinical trials using the available clinical grade FUS-MB and BPN at our and other institutions. The approaches can also be easily adapted for use in other cancers as well.

## Materials and Methods

### Computational and experimental workflow for identifying putative master regulatory miRNAs

Our new methodology for identifying key regulatory microRNAs (miRNAs) and their prioritization involved several steps. First, all miRNA mRNA targets were identified through PAR-CLIP **(described in more detail in the supplementary methods)**. Argonaute/target gene complexes were collected, the argonaute protein was digested, and the complexed target genes were then sequenced. T/C alignment analysis was used to identify genes with the distinctive mutation that identifies genes that complex with miRNAs. The sequence target fragments were then compared with a list of known miRNA seed sequences using string sequencing matching in R. Specifically, miRNAs that targeted genes in the 3’-untranslated region were prioritized, while genes with no miRNA matches or matches in the coding or 5’-untranslated regions were filtered out.

Next, we analyzed 166 Glioblastoma Multiforme (GBM) RNA-Seq datasets from The Cancer Genome Atlas (TCGA), which were normalized against 255 normal brain datasets from the Genotype-Tissue Expression (GTEx, n = 255) project using the bowtie and bedtools genomic sequencing alignment packages. For 42,644 genes, these tumor samples were compared to normal using the DESeq2 package in R, revealing genes that were greater than 2-fold up- and downregulated. Prior to downstream analyses, we implemented pre-filtering measures, discarding genes with read counts of less than five reads.

In the third step, the CancerMine database was employed to classify each miRNA target gene as an oncogene (ONC), a tumor suppressor gene (TSG), or neither, using experimentally verified labels from published manuscripts. Using this method, the final curated repository contained 1849 published oncogenes and 1478 published tumor suppressors, a 1.25:1 ratio.

The fourth step involved performing a survival analysis on the significantly deregulated genes from the previous differential expression analysis. Using the R programming package, survminer, a cox-proportional hazard ratio was determined for each putative target gene.

The fifth step involved filtering out targets that did not have a “consistent” and “significant” survival and deregulation trend. For oncogenic targets, these parameters were determined as a minimum 2-fold increase in expression and a multiple hypothesis adjusted p-value of ≤0.05, coupled with either an inconclusive correlation with survival (cox coefficient between −0.3 and 0.3) or a correlation with poor prognosis (cox coefficient >0.3). Tumor suppressive targets required a minimum 2-fold decrease in expression, an adjusted p-value of ≤0.05, and either an inconclusive correlation with survival or a correlation with good prognosis (cox coefficient <-0.3). Targets that met these criteria were labeled “consistent” and were retained for the next step.

For this sixth step, the retained “consistent” targets were assigned a composite score based on the percentile ranks of their absolute expression and differential expression scores. For example, a target at the 100th percentile would score 1, and at the 0th percentile would score 0, with the highest possible score being 2 for targets at the 100th percentile for both absolute and differential expression.

Subsequently, each miRNA received two scores reflecting the composite scores of their annotated targets: a TSG score from oncogenic targets and an ONC score from tumor suppressive targets. These scores were then ranked by percentile.

In the final analysis, the percentile rank difference between the TSG and ONC scores for each <u>miRNA</u> determined its classification. The ONC percentile rank was subtracted from the TSG rank. A difference exceeding 25% classified a miRNA as a TSG miRNA (TSG>ONC), while a difference below −25% classified it as an ONC miRNA (ONC>TSG). This threshold was derived from the 1.25:1 oncogene enrichment in our Cancermine dataset described in step three of this analysis. For instance, miR-1185, with a TSG score in the 84th percentile and an ONC score in the 12th percentile, was identified as a projected tumor suppressor miRNA (+72% difference). This approach also minimized the emphasis on miRNAs targeting numerous ONC and TSG genes, as they would rank highly in both categories. For example, miR-4709 ranked in the 100th percentile for TSG and 90th percentile for ONC, but would receive “inconclusive” classification by our algorithm, as there is only a 10% score difference in these roles.

### Photoactivatable-ribonucleoside-enhanced crosslinking and Immunoprecipitation (PAR-CLIP)

A total of 1 µg of FLAG-tagged AGO1, AGO2, AGO3 plasmids were transfected into 2X10^5^ U87 cells via Lipofectamine 2000 transfection. Following transfection, the cells were subjected to puromycin selection at a concentration of 1 µg/ml for 48 hours to generate stable cell lines. Verification of AGO1, AGO2, AGO3 overexpression was conducted through immunoblot analysis employing AGO1, AGO2, AGO3 antibodies. The PAR-CLIP methodology was implemented and adapted from the previously established protocol (*33*). Briefly, AGO1, AGO2, AGO3 stable cell lines were cultured overnight in the presence of 100 µM 4-thiouracil (4SU), to label all cellular nascent RNA. Subsequently, these 4SU-labeled cells were exposed to 365 nm UV light (utilizing a Spectro Linker XL-1500) to effectuate the crosslinking of labeled nascent RNA with RNA binding proteins. Following UV exposure, cells were rinsed twice with 1X PBS, harvested, and lysed using a buffer composed of 50 mM HEPES-KOH (pH 7.5), 150 mM KCl, two mM EDTA (pH 8.0), one mM NaF, 0.5% NP-40, 0.5 mM DTT, and freshly prepared protease inhibitor cocktails. Cell lysates were centrifuged at 15,000Xg for 15 minutes at 4^0^C. The supernatant from the lysed cells was subjected to treatment with RNAse T1 at a concentration of 1 U/µl for 15 minutes at room temperature. Subsequently, the Flag-tagged AGO1, AGO2, AGO3 cell lysates’ supernatant underwent immunoprecipitation, facilitated by an anti-Flag antibody conjugated to Protein G Dynabeads. The immunoprecipitated material was further subjected to digestion with RNase T1, and the resulting beads were subjected to washing with a high salt wash buffer. The washed beads were re-suspended in a dephosphorylation buffer and treated with calf intestinal alkaline phosphatase for 10 minutes at 37^0^C to eliminate phosphate groups from RNA molecules. Next, the dephosphorylated beads were treated with polynucleotide kinase and radioactive [γ-32P]-ATP for 30 minutes at 37^0^C to introduce RNA labeling. The protein-RNA complexes were then separated via SDS-PAGE and subsequently electroeluted. These complexes exhibited a migration pattern at around 100 kDa. The electroeluted samples then underwent digestion with proteinase K to release RNA from the complexes, followed by RNA extraction involving an acid phenol/chloroform mixture and ethanol precipitation. The extracted RNA was then converted into cDNA, and adaptor ligation was executed following previously established protocols (*31, 65*). The resulting libraries were sequenced, and the generated short reads were mapped against the human genome hg38, mRNA, and miRNA precursor databases and the clustered sequences were identified. The PAR-CLIP clustered sequences were identified by T to C conversion at the 4-thiouridine cross-linked site. The majority of the clustered sequences were found at the 3’-UTR regions. Finally, the true target sites were determined from the clustered sequences based on the list of input miRNA seed sequences.

### DNA-Brain penetrating nanocomplexes (BPNs) synthesis and characterization

The preparation of BPN formulation was conducted in accordance with the protocol outlined in a previous publication (Mastorakos et al., 2017). In summary, poly(beta-amino esters) (PBAE) and PEGylated PBAE (PEG-PBAE) were synthesized through the conjugation of 1,11-diamino-3,6,9-trioxaundecane (obtained from Millipore Sigma, St. Louis, MO) or 5 kDa methoxy-PEG-N-hydrosuccinimide (mPEG-NHS, 5 kDa, from Sigma-Aldrich) with the acrylate groups on PBAE (sourced from Sigma-Aldrich), respectively. A mixture of PBAE and PEG-PBAE polymers was then prepared at a 3:2 ratio by PBAE amount to create a highly PEGylated surface. For the formulation of DNA-BPN, polymers and nucleic acids were vigorously mixed at a weight ratio of 60 and a volume ratio of 1:5. This mixture was allowed to incubate at room temperature for 30 minutes to facilitate nanoparticle (NP) assembly. Subsequently, the solution was placed into 100 kDa MWCO Amicon Ultra Centrifugal Filters (Millipore Sigma) and centrifuged at 1,000 × g for 15 minutes at 4°C. To remove residual polymers from the DNA-BPN solution, the concentrated DNA-BPNs, with a nucleic acid concentration adjusted to 1 mg/mL, were diluted tenfold with DNase/RNase-Free Distilled Water and re-centrifuged under the same conditions. After undergoing two additional washing steps, the final DNA-BPN solution, with a nucleic acid concentration of 1 mg/mL, was prepared for subsequent in vivo experiments.

### MRI-guided Focused Ultrasound (MRIgFUS)

All aspects of animal handling, monitoring, housing, and experiments strictly adhered to the guidelines set forth by the National Institutes of Health and complied with local Institutional Animal Care and Use Committee regulations at the University of Virginia. To establish tumors, immunodeficient mice aged 6-8 weeks were utilized. Specifically, 3X10^5^ U87 cells were intracranially introduced into the striata of immunodeficient mice. After a three-week injection period, MRI-guided focused ultrasound (MRIgFUS) was employed following a well-established protocol, with some modifications(*70*). Briefly, the MRIgFUS experimental setup featured an MRI-compatible pre-focused eight-element phased array, accompanied by a 1.5 MHz geometrically focused transducer boasting a 25 mm active diameter and a focal ratio of 0.8. These components were interconnected through a phased array generator and a radiofrequency power amplifier. It was connected to an MRI-compatible motorized stage to precisely control the transducer’s movements in the rostral-caudal and medial-lateral orientations. Degassed water was introduced into the spherical transducer’s membrane to ensure effective coupling between the membrane and the mice’s brains. At the same time, acoustic gel was applied to both the inflated membrane and the shaved portion of the mice’s skull. These measures prevented the entrapment of air bubbles. For intravenous injections, a catheter was inserted into the mouse’s tail. Microbubbles (25 µl/kg body weight) and an MRI contrasting agent, gadobenate dimeglumine (0.1 ml), were administered via this catheter with saline. Subsequently, a series of MRI images were captured. The precise positioning of the mouse’s brain relative to the transducer was determined by locating the transducer’s position within the MRI space. During sonication and MRI imaging, the mice were positioned in a prone posture. The region of interest (ROI) encompassing the tumor was defined, and sonication was carried out using a 1.5 MHz transducer to induce the opening of the blood-brain barrier around the tumor. MRI images were acquired using a surface coil incorporated into the FUS system. The effectiveness of blood-brain barrier opening was verified by comparing MRI sections before and after sonication. This MRI-guided focused ultrasound approach provided a powerful method for non-invasive modulation of the blood-brain barrier, enabling targeted delivery of therapeutic agents to brain tumors in preclinical models.

### Detailed descriptions of all other methods can be found in the supplementary material

### Statistical analyses

All the data are represented as mean ± S.E.M. from three independent biological replicates. The P value is calculated by two tailed Student’s t-tests when comparing two groups. P values of < 0.05 were considered statistically significant.

## Acknowledgements

We thank Drs. Jeongwu Lee, Cleveland Clinic and Erik P. Sulman, NYU Langone Health for providing the glioblastoma stem cell lines GSC-28 and GSC-34 respectively. This work is supported by the University of Virginia Bioinformatics Core, Molecular Imaging Core, Advanced Microscopy Facility and Research Histology Core. We would also like to thank dbGAP and TCGA data management team for providing access to the raw RNA-seq data.

## Author Contributions

Conceptualization: SS, YZ, MKG, CD, RA

Methodology: SS, YZ, MKG, CD, FH, EQXM, PM, XW, GK, NC, RRC, FG, PK, ALK, JSS, JM

Investigation: SS, YZ, MKG, CD, RA

Visualization: SS, YZ, CD, MKG, FH, PM, MD

Funding acquisition: RA

Project administration: RA Supervision: RA

Writing – original draft: SS and RA

Writing – review & editing: all the authors

## Data and materials availability

The PAR-CLIP dataset generated in this study has been deposited in the Gene Expression Omnibus (GEO) under accession number GSE293517. All codes and materials supporting the findings of this study are available from the corresponding author upon reasonable request.

## Supplementary methods

### Cell culture and transfection

All glioblastoma cell lines used in this study were procured from the American Type Culture Collection (ATCC). Specifically, U87 cells were cultured and maintained in Minimal Essential Medium (MEM) supplemented with one mM sodium pyruvate, one mM non-essential amino acids (NEAA), 10 ml of 7.5% sodium bicarbonate, 1% penicillin/streptomycin, and 10% fetal bovine serum. A172 and 293T cells were cultured in DMEM high glucose medium supplemented with 1% penicillin/streptomycin and 10% fetal bovine serum. Glioblastoma stem cells, GSC-34, we from Cleveland Clinic, and GSC-28 were obtained from MD Anderson Cancer Center. These stem cells were cultured following a previously published protocol (*25*). The culture medium for stem cells consisted of 2.5 ml of N-2 (0.5X), 5 ml of B27 (0.5X), 50 ng/ml EGF, 50 ng/ml FGF, 0.5 mM glutamine, and 1% penicillin/streptomycin. Regular maintenance included dissociation using an accutase solution when the neurospheres reached a specific size. The primary astrocytes were from Lonza Bisoscience and were cultured in growth medium kit from Lonza Bioscience. All cells were cultured at 37°C in a humidified incubator with a 5% CO2 atmosphere to maintain cell viability. Both glioblastoma and GSCs were transiently transfected with either 30 nM of a scrambled negative control miRNA, target-specific mimic primary miRNA, or 30-50 nM of a miRNA inhibitor, using the RNAiMax reagent. Following transfection, the culture medium was replaced with a complete growth medium after 6-8 hours. Subsequently, cells were harvested 48 hours post-transfection for various analyses, including RNA isolation, cDNA synthesis, qPCR, and cell lysate preparation for immunoblot analysis to validate miRNA targets.

### 3’UTR reporter assay

We conducted luciferase reporter assays to investigate miRNA interactions with specific target genes according to the previously published literature (*25*). First, we PCR amplified the 3’-UTRs (3’ untranslated regions) targeted by miR-340, miR-382 and miR-17 from genomic DNA. These amplified sequences were then cloned into the psiCheck2 vector, which had been digested with BamH1 and Not1 restriction enzymes. The primers used for amplifying genomic DNA are shown in **Supplementary Table 1**. All reporter plasmids were subjected to Sanger sequencing and verification to ensure sequence accuracy. for the reporter assay, we co-transfected the primary miRNAs along with 1 μg of the psiCheck2 reporter plasmid into our cells of interest. After 48 hours of transfection, we lysed the cells using 1X passive lysis buffer. Subsequently, we performed a luciferase reporter assay using a Promega dual luciferase reporter assay kit. Firefly luciferase activity served as an internal control to normalize the luciferase signal, allowing us to assess the interaction of miRNAs on the regulation of specific target genes.

### Invasion assay

To evaluate the effects of miRNAs on cellular invasion, we followed a previously published protocol (*66*). In brief, transwell invasion chambers were coated with 75 μl of a 250 μg/ml collagen IV solution. After 48 hours of miRNA transfection, cells were counted, and 1 × 10^5^ cells were re-suspended in 300 μl of 1% serum-containing, antibiotic-free medium. The cells were then added to each transwell chamber, with 750 μl of fetal bovine serum-containing medium in the bottom chamber, creating a gradient that stimulates cell invasion through the collagen IV-coated membrane. Following 6-8 hours of incubation, the invaded cells were stained with crystal violet, and five random fields were captured and counted using ImageJ software.

### Gel electrophoresis and immunoblot analysis

Immunoblot assay was performed according to previously published protocols (*67*). After 48 hours of miRNA transfection, cells were harvested and cell lysis was performed using RIPA buffer (50 mM Tris-HCl pH 8.0, 150 mM NaCl, 5 mM EDTA, 0.1% SDS, and 1X protease and phosphatase inhibitor). The protein concentration in the lysates was determined using the Bradford method. Subsequently, 40-100 µg of cell lysates were loaded onto each lane, separated via 4-12% SDS-PAGE, and transferred to a nitrocellulose membrane. To prevent non-specific binding, the membrane was blocked with 5% skim milk and then incubated overnight at 4^0^C with primary antibodies targeting specific proteins, including CD44 (1:1000, Catalog#37259,Cell Signaling Technology), TOP2A(1:500, Catalog#sc-3659, Santa Cruz Biotechnology), ANKRD11(1:1000, Catalog# sc-81049, Santa Cruz Biotechnology), ZBTB4(1:1000, Catalog#sc-514883, Santa Cruz Biotechnology), EPHA4(1:1000, Catalog#sc-365503, Santa Cruz Biotechnology), EHD3(1:1000, Catalog#sc-100723, Santa Cruz Biotechnology), GAPDH(1:2000, Catalog#sc-166545, Santa Cruz Biotechnology), MDM2(1:1000, Catalog#OP115, Sigma Aldrich), RHOC(1:1000, Catalog# 10632-1-AP, Thermo Fisher Scientific), HMGA2(1:1000, Catalog#8179, Cell Signaling Technology), PLAU(1:1000, Catalog# 17968-1-AP, Thermo Fisher Scientific), NUSAP1(1:1000, Catalog#H00051203-B01P). After primary antibody incubation, the membrane was probed with horseradish peroxidase-conjugated secondary antibodies (goat anti-mouse or rabbit) at room temperature for 1 hour. Detection was achieved using a pico-sensitive ECL reagent (Thermo Scientific), and images were captured using an Amersham 600 imager.

### Cell proliferation assay

2X10^4^-5X10^4^ U87 and U251 cells were seeded and transfected with 30 nM of either scrambled negative control miRNA or primary mimic miR-340, miR-382, or miR-17 inhibitor. After a transfection period of 48 hours, the cells were trypsinized and counted at various time points for seven days. The resulting cell counts were then used to generate growth curves.

### RNA isolation, cDNA synthesis, and qPCR

Total RNA was extracted from primary astrocytes, U87, A172, U251, GSC-28 and GSC-34 cells using the Qiagen miRNeasy kit following the manufacturer’s protocol. GBM tissue samples were homogenized, and total RNA was isolated using the same kit. Subsequently, 100 ng to 1 µg of total RNA was employed for cDNA synthesis utilizing either the miRscript or miRCURY LNA RT kit. The miRCURY LNA SYBR Green PCR kit was used to assess the expression levels of mature miRNAs. The miRNA fold change was calculated using the delta-delta Ct method, with primary astrocytes used as a control sample to normalize the fold change.

### Neurosphere assay

The neurosphere assay was conducted according to a previously published protocol (*66*). Briefly a 6-well plate was initially coated with a poly-ornithine solution and incubated at 37^0^C in a tissue culture incubator for one hour. Afterward, the plates were washed twice with 1X PBS. Glioblastoma stem cells GSC-34 and GSC-28 were dissociated and seeded onto the coated plate. Transfection with the scrambled negative control and the primary mimic miR-340, miR-382, and inhibitor against miR-17 was conducted using the Lipofectamine RNAiMax reagent. After 48 hours of transfection, glioblastoma stem cells were dissociated using an accutase solution and cultured in a neurobasal growth medium for another seven days. The number of neurospheres was quantified both manually and also with ImageJ software based on their size classification (large, medium, and small) and represented in a bar graph.

### In vivo effects of miRNA expression

To assess the impact of miRNA overexpression or inhibition in vivo, a mouse xenograft model was employed. Six-week-old athymic nude mice were obtained from the Jackson Laboratory. U87 cells (3X10^5^) were seeded in a 6-well plate and transfected with 30 nM of either scrambled negative control miRNA or precursor mir-340, miR-382, or mir-17 inhibitor. After 48 hours of transfection, the cells were trypsinized, counted, and utilized for intracranial injection. All mouse experiments were conducted following the University of Virginia Institutional guidelines. The transfected cell (3X10^5^) were intracranially injected and stereotactically implanted into the striata of immunodeficient mice, according to a previously established protocol (*66, 67*). After three weeks, MRI imaging was used to visualize and quantify tumor growth. To enhance tumor signal intensity, gadopentetate dimeglumine was intraperitoneally injected. Imaging was conducted using T1-weighted coronal sections with a field size of 2.5 mm × 2.5 mm and a resolution of 256×256 pixels. Osirix lite software was used to calculate tumor volume using MRI tumor images.

### Microbubbles preparation

Gas microbubbles were produced in-house using a method previously described (*68*) involving sonication of decafluorobutane gas. The process begins with distearoyl phosphatidylcholine (Lipoid, Germany, or Avanti Polar Lipids, Alabama) and PEG150 monostearate (obtained from Stepan Kessco, Illinois) being dispersed in normal saline at a concentration of 2 mg/ml for each component. This mixture is then sonicated at 20 KHz and 30% power using a Q700 Sonicator (Qsonica, Newton, CT) for 3 minutes to ensure the formation of a submicron micellar dispersion.

To introduce decafluorobutane into the mixture, the headspace of a vial was filled with the gas via a teflon capillary tubing that sparged the gas through the aqueous medium. Sonication was then continued at maximum power for an additional 30 seconds to disperse the gas into the medium and form microbubbles. Following sonication, the sample was allowed to cool and then floated at normal gravity to remove any bubbles larger than 5 µm, in accordance with the guidelines similar to those in {Klibanov, 2004 #104}. The prepared microbubbles were then dispensed into individual vials under a decafluorobutane atmosphere and the vials were stoppered and stored under refrigeration, ensuring they were not frozen. These gas microbubbles are characterized by their neutral nature.

## Supplementary Figure Legends

**Figure S1:**
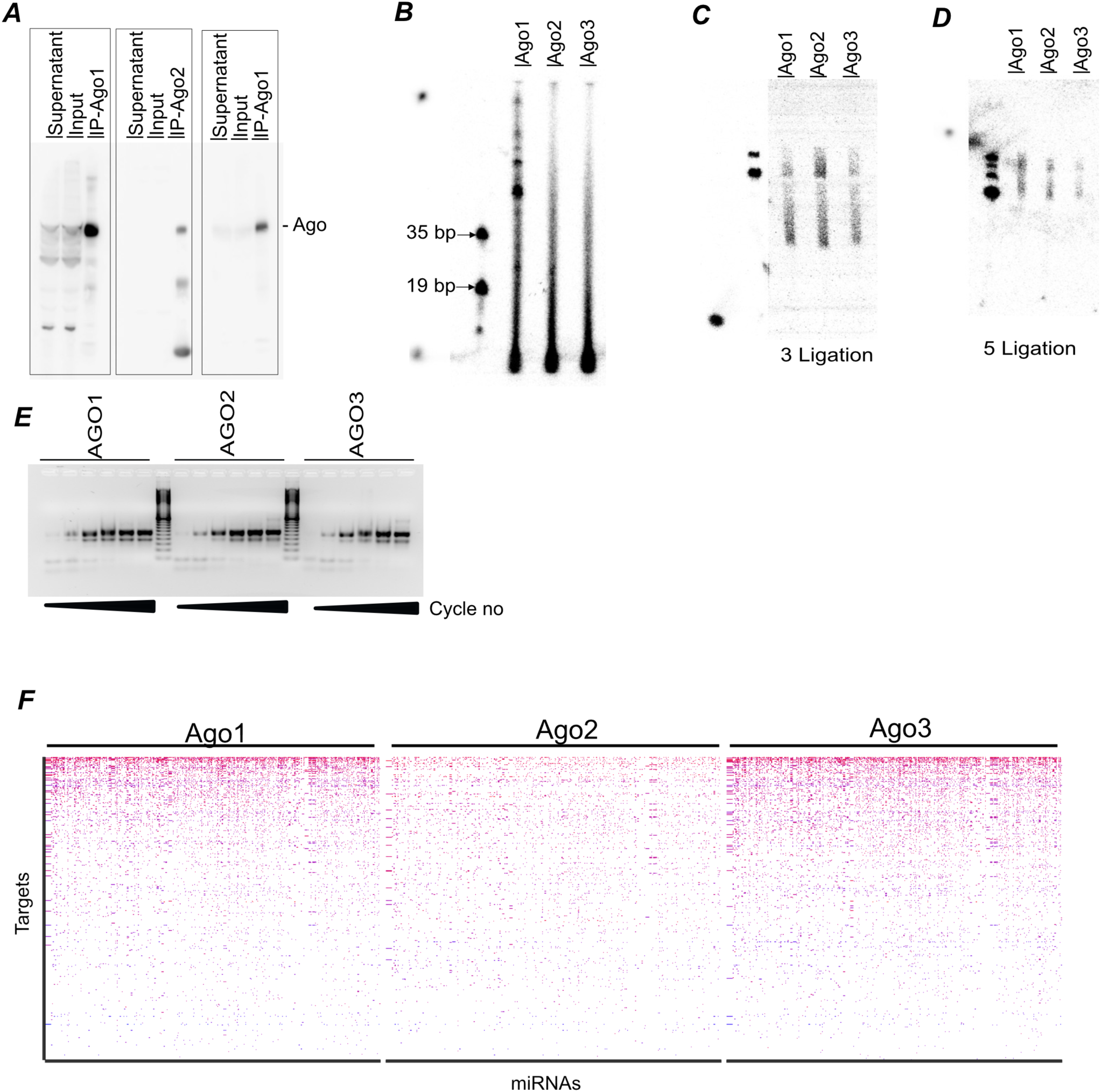
Identification of miRNA targets through PAR-CLIP in glioblastoma cells: A) U87 cells stably expressing Flag-tagged AGO1, AGO2, AGO3 were immunoprecipitated with Flag antibody and immunoblotted with Flag antibody to detect Ago1, Ago2, and Ago3. Supernatant and input were examined to assess pull-down efficiency. The bands at the labeled AGO position show successful pulldown of Ago1, Ago2 and Ago3 respectfully. B) Radiolabeled RNA fragments purified from AGO1, AGO2, AGO3-RNA immuno-complexes were separated on a 15% polyacrylamide gel, and regions of 19-35 bp were excised and purified for 3’ and 5’ adaptor ligation reactions. C, D) 3’ (C) and 5’ (D) adaptor ligation reactions were conducted as described in the materials and methods section which shown successful incorporation of adaptors. E) The ligated RNA fragments were reverse transcribed, and libraries were generated for AGO1, AGO2, AGO3 samples. The libraries from AGO1, AGO2, AGO3 were subjected to amplification at different cycles to prevent saturation. F) Heatmaps displaying all the miRNA targets identified in U87 cells by AGO1, AGO2, AGO3 through PAR-CLIP which show that each miRNA exhibits numerous targets across the different AGOs.

**Figure S2:**
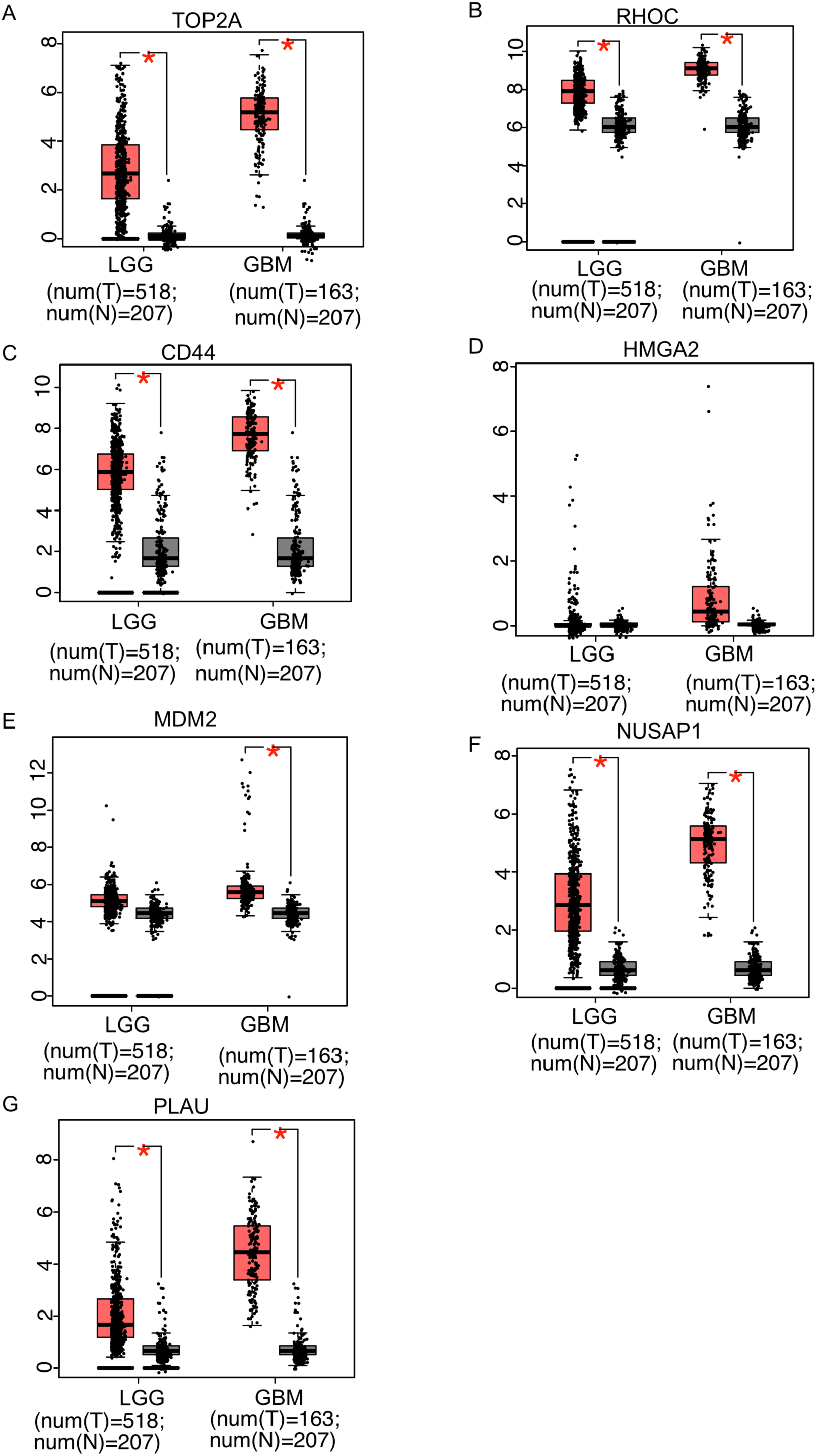
Expression of miR-340 and miR-382 targets in TCGA glioma and GTEx normal brain samples. A–G) Expression levels of validated miR-340 and miR-382 targets — *TOP2A*, *CD44*, *RHOC*, *MDM2*, *HMGA2*, *NUSAP1*, and *PLAU* — were analyzed using the GEPIA online tool in TCGA glioma and GTEx normal brain samples. All targets were significantly upregulated in tumor samples (red bars) compared to normal samples (gray bars). *P* < 0.05.

**Figure S3:**
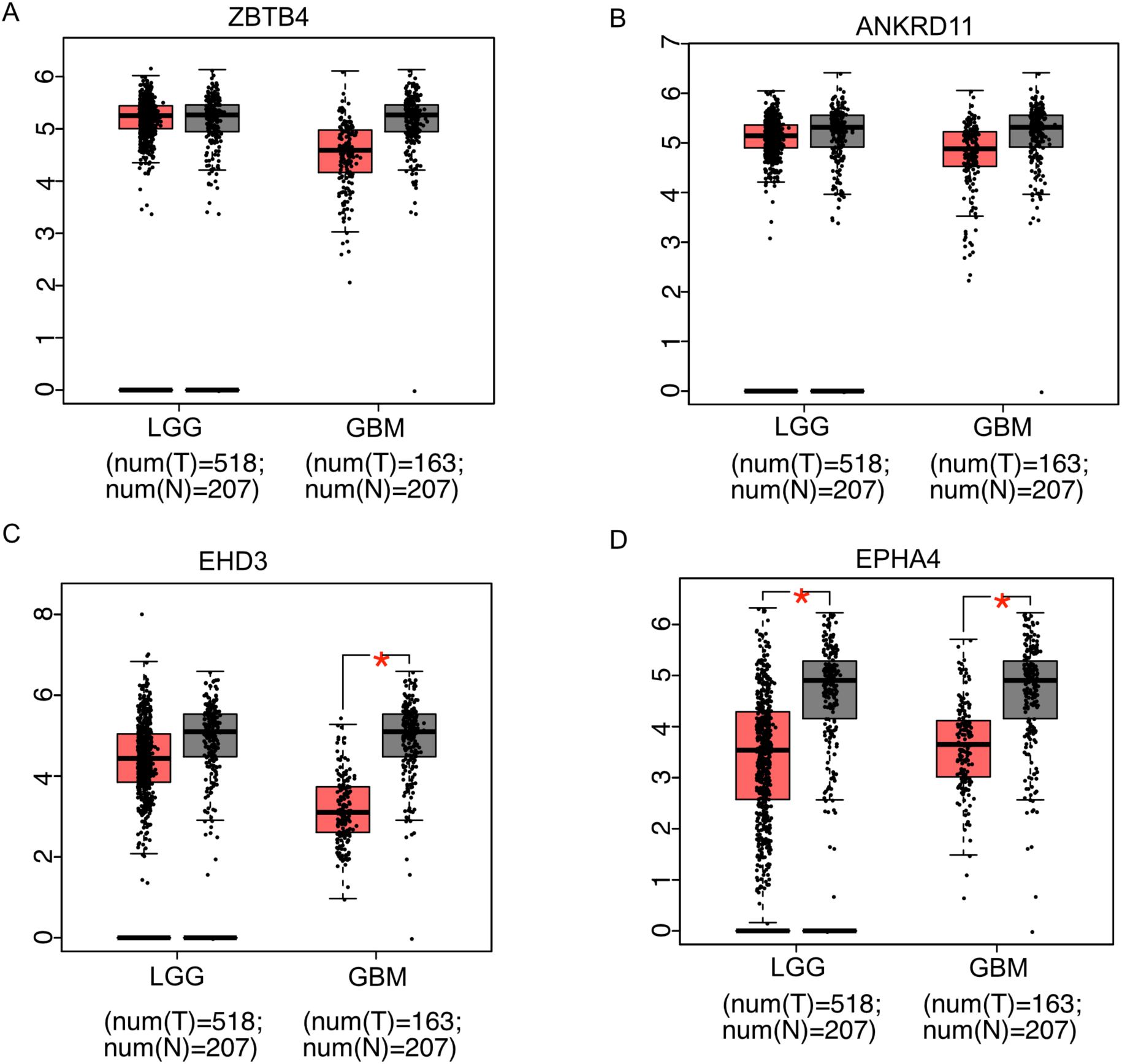
Expression of miR-17 targets in TCGA glioma and GTEx normal brain samples. A–D) Expression analysis of miR-17 targets — *ZBTB4*, *ANKRD11*, *EHD3*, and *EPHA4* — using GEPIA shows significant downregulation in glioma samples relative to normal brain tissues. *P* < 0.05.

**Figure S4:**
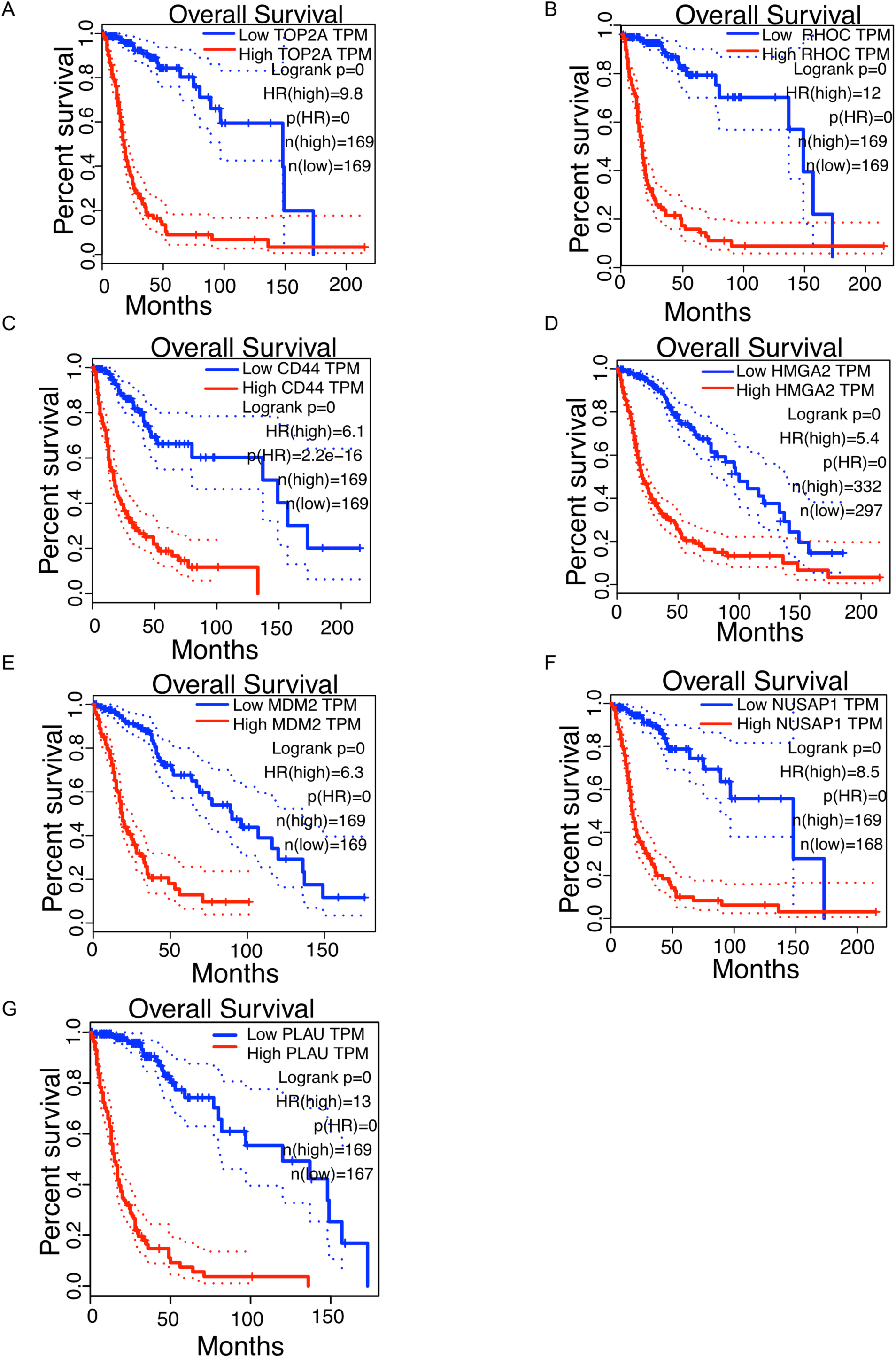
Kaplan–Meier survival analysis of tumor-suppressive miR-340 and miR-382 targets in TCGA primary glioma patients. A–G) Survival analysis was conducted using the GEPIA tool based on TCGA glioma cohorts. Patients were stratified into high and low expression groups for each target gene. High expression of the target gene (red line) was significantly associated with poor prognosis. *P* < 0.05.

**Figure S5:**
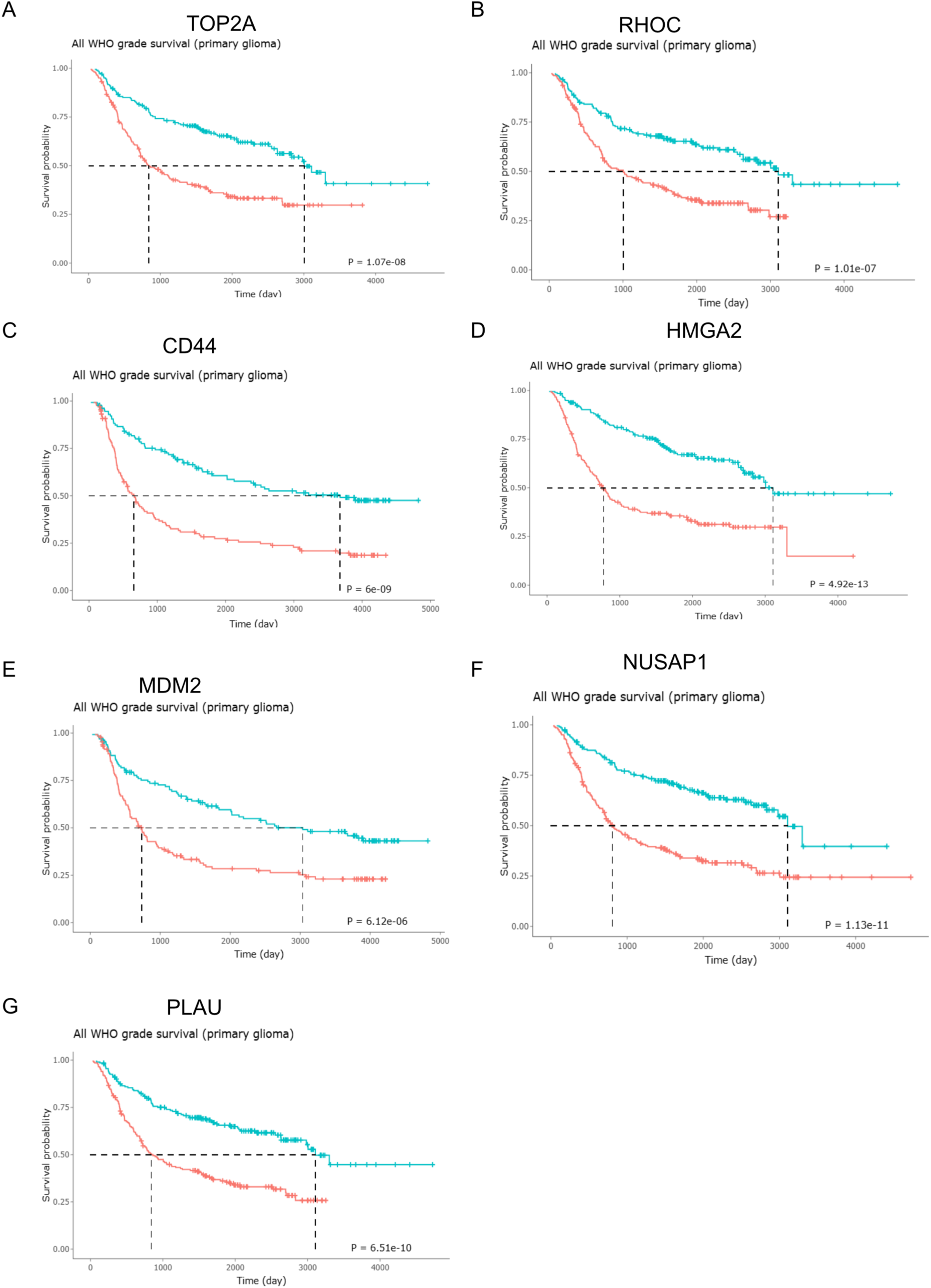
Kaplan–Meier survival analysis of miR-340 and miR-382 targets in CGGA primary glioma patients. A–G) Using the CGGA glioma dataset, high expression of the validated tumor-suppressive targets of miR-340 and miR-382 (red line) was significantly associated with poor prognosis. *P* < 0.05.

**Figure S6:**
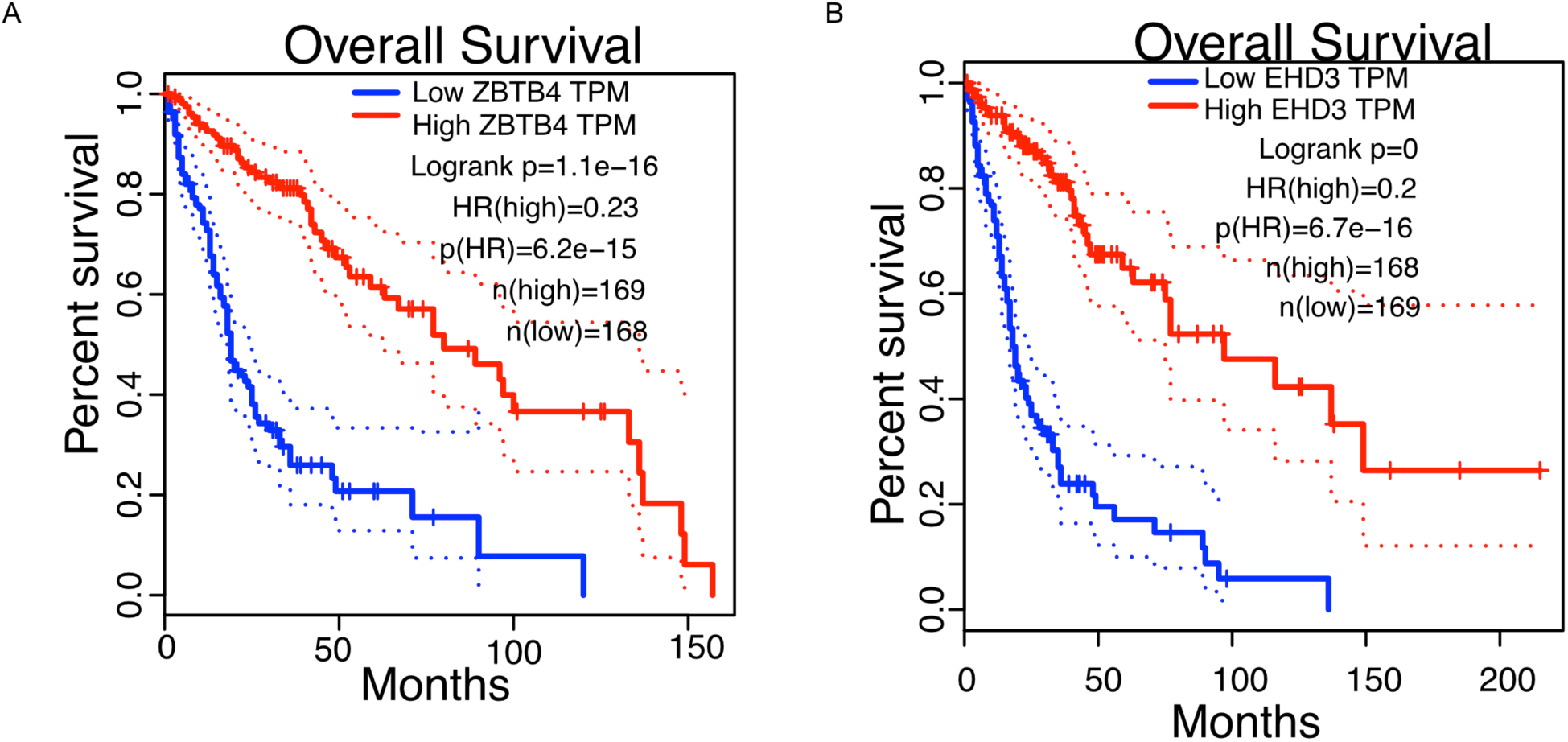
Kaplan–Meier survival analysis of oncogenic miR-17 targets in TCGA primary glioma patients. A,B) Survival analysis of selected miR-17 targets in TCGA glioma cohorts. High expression of the target gene (red line) was significantly associated with better prognosis. *P* < 0.05.

**Figure S7:**
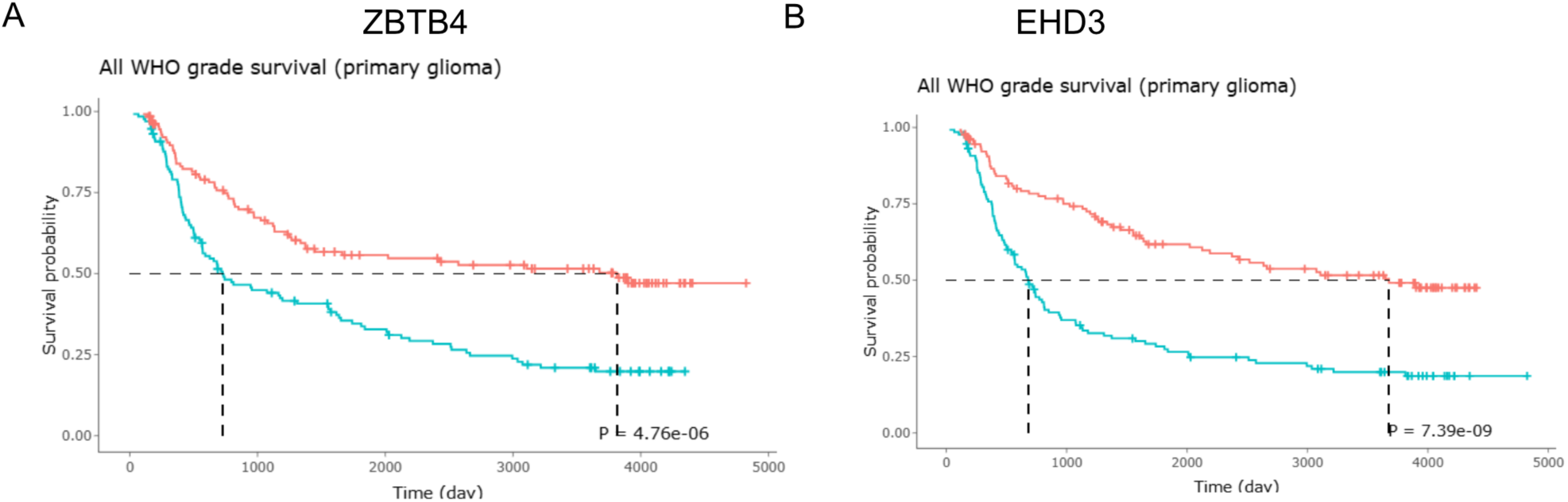
Kaplan–Meier survival analysis of oncogenic miR-17 targets in CGGA primary glioma patients. A,B) Kaplan–Meier analysis of selected miR-17 targets in CGGA glioma samples showed that high expression (red line) was associated with improved survival. *P* < 0.05.

**Figure S8:**
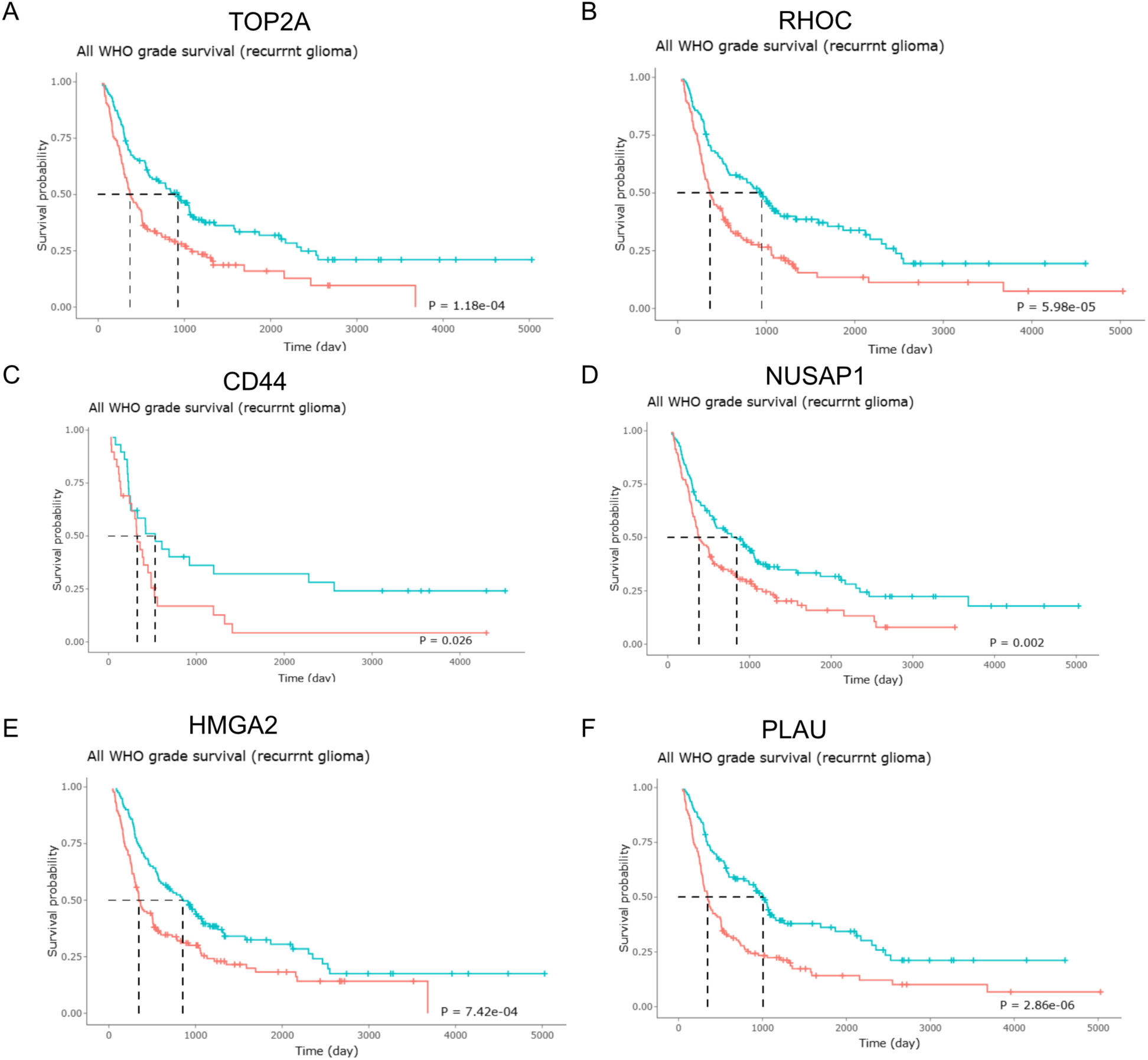
Kaplan–Meier survival analysis of miR-340 and miR-382 targets in CGGA primary glioma recurrent patients. (A–G) High expression of validated tumor-suppressive targets of miR-340 and miR-382 (red lines) was significantly associated with poor prognosis in the CGGA recurrent glioma dataset (*P* < 0.05).

**Figure S9.**
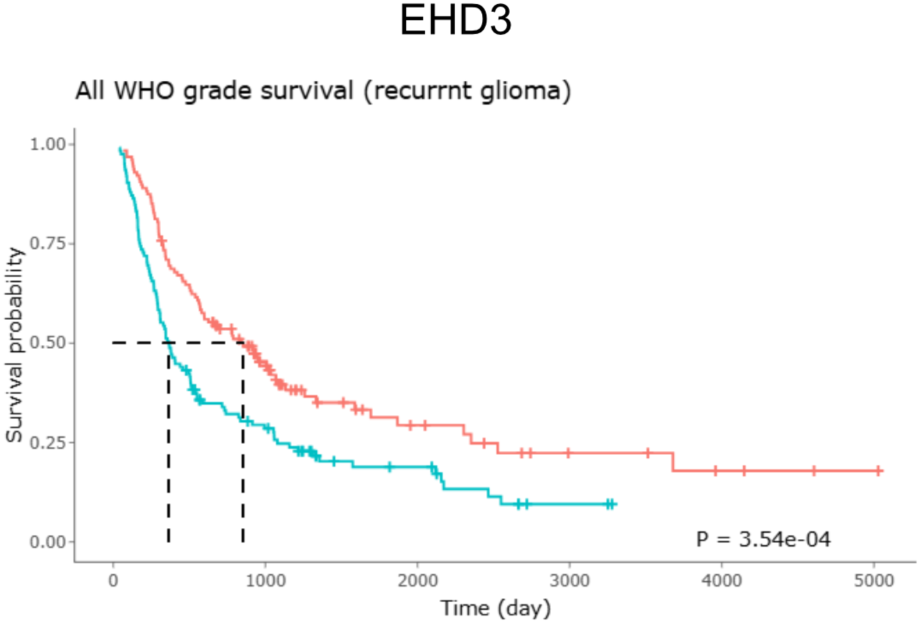
Kaplan–Meier survival analysis of the oncogenic miR-17 target EHD3 in recurrent glioma patients from the CGGA primary glioma dataset. Patients were stratified into high and low EHD3 expression groups using the CGGA online analysis tool. High EHD3 expression was significantly associated with improved overall survival in recurrent glioma (*P* < 0.05).

**Figure S10:**
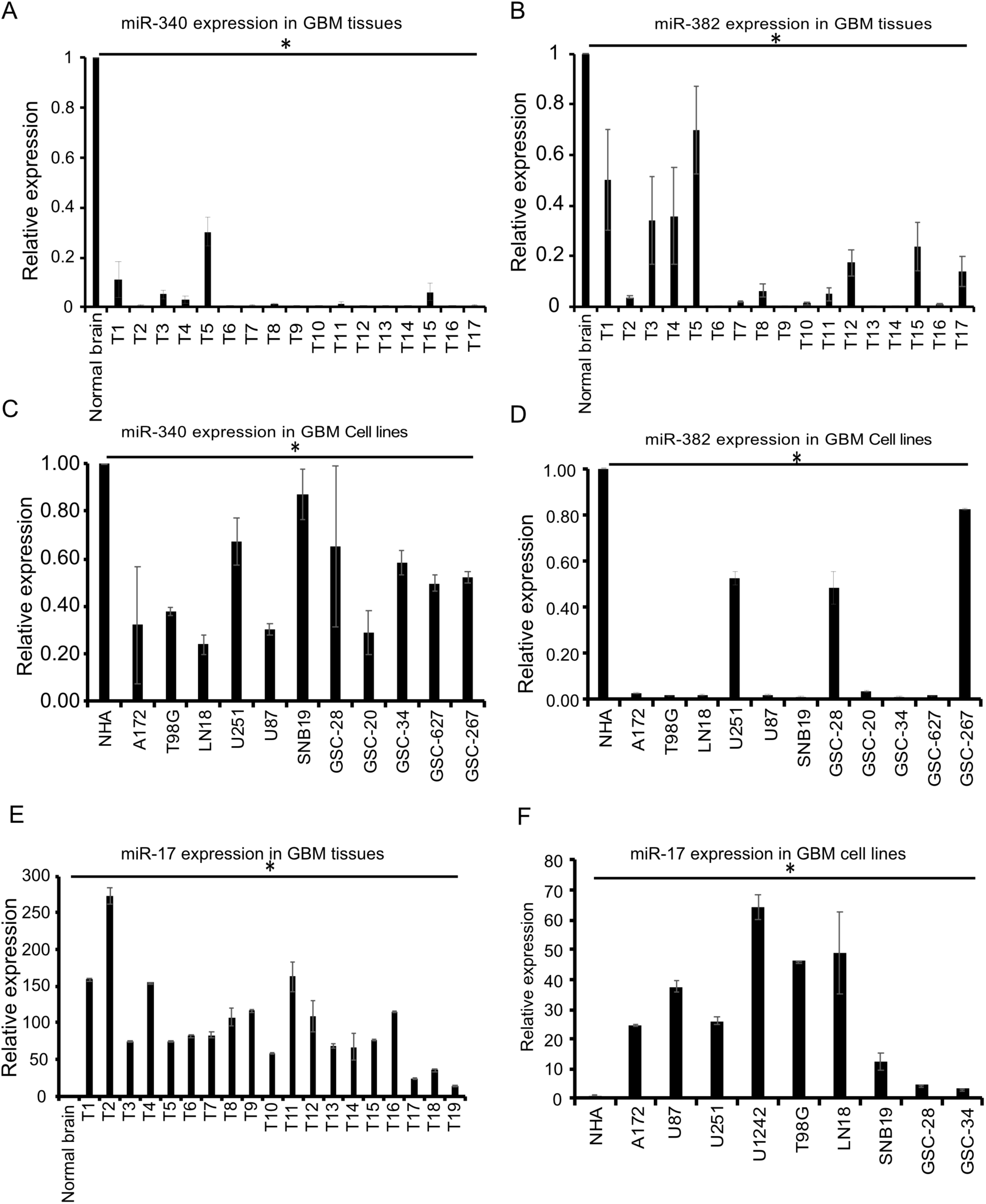
Endogenous expression of miR-340, miR-382, and miR-17 in different glioblastoma cell lines and tissues: A,B) Total RNA was extracted from various glioblastoma tissue samples, followed by cDNA synthesis using the Qiagen miScript RT kit. One microgram of RNA was utilized for cDNA synthesis. The cDNA samples were diluted, and RT-qPCR was conducted with miRNA primers against miR-340 (A), miR-382 (B). Normal brain samples were used as controls, and U6 was employed as an internal control for normalization. MiR-340 and miR-382 showed decreased expression compared to normal brain samples. C,D) Different glioblastoma cell lines, including A172, T98G, LN18, U251, U87, SNB19, and patient-derived glioblastoma cell lines (GSC-28, GSC-20, GSC-34, GSC-627, GSC-267) were cultured, and cells were harvested for total RNA isolation using the Qiagen miRNeasy kit. Normal human astrocytes (NHA) were used as normal controls. One microgram of total RNA was used for cDNA synthesis with the miRCURY kit, followed by qPCR with primers against miR-340 (C), miR-382 (D). MiR-340 and miR-382 showed decreased expression compared to normal human astrocytes. E) Total RNA was extracted from different glioblastoma tissue samples followed by cDNA synthesis with miScript RT kit using miR-17 primer set. MiR-17 showed elevated expression in all the glioblastoma tissue samples. F) Expression of miR-17 in various glioma cell lines and patient derived glioma stem cell lines by qPCR. miR-17 showed elevated expressed compared to normal human astrocytes. Results are from three independent experiments. * P < 0.05.

**Figure S11:**
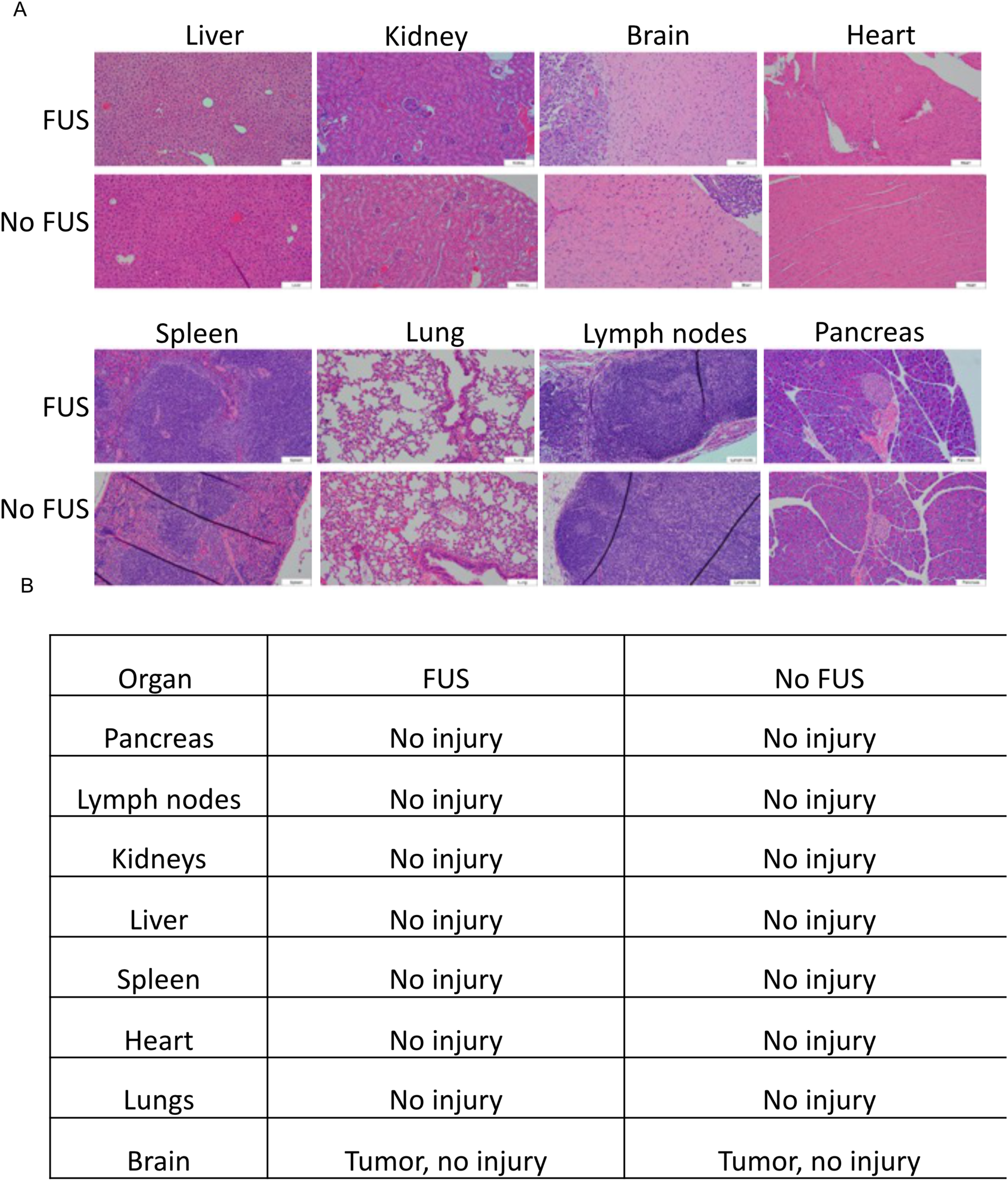
Magnetic Resonance-guided Focused Ultrasound (MRgFUS)– microbubbles and brain penetrating nanoparticle (FUS-MB-BPN) mediated delivery of miRNA into mouse brain tumors does not induce toxicity. A) Panel A illustrates the delivery of BPN-conjugated miR-142-3p into the mouse brain using MRgFUS, followed by harvesting of various organs under both FUS and non-FUS conditions, with subsequent H&E staining. B) Panel B presents a table summarizing toxicity assessments by an experienced neuropathologist across different organs Liver, Kidney, Brain, Heart, Spleen, Lung, Lymph nodes, Pancreas which showed there was no apparent off-target toxicity to the organs.

